# Delayed bleaching-induced coral mortality limits reef recovery after a marine heatwave

**DOI:** 10.64898/2026.10.07.757404

**Authors:** Andreas Eich, Yannick Chancerelle, Valeriano Parravicini, Nicolas Derome, Laetitia Hédouin, Caroline E Dubé

**Author notes:** **Corresponding author:** Andreas Eich. Should be considered joint senior authors (Laetitia Hédouin, Caroline E Dubé).

## Abstract

Coral reefs worldwide are increasingly exposed to marine heatwaves, which cause recurrent coral bleaching and mass mortality events. While the immediate impact of bleaching has been extensively studied, the longer-term consequences and recovery trajectories remain poorly understood. By integrating two long-term monitoring datasets, we quantified how fine-scale variation in heat stress exposure shaped post-bleaching trajectories. We examined patterns of coral mortality and recovery across 13 sites in Mo’orea (French Polynesia) after the most severe marine heatwave recorded on the island in 2019. The maximum accumulated heat stress (AHS_max_) varied between sites (from 5.76 to 6.26 °C weeks), allowing the assessment of heat stress-related mortality between coral genera. *Pocillopora* mortality increased with AHS_max_, and mean coral cover collapsed from 35% to 17% within one year. *Acropora* experienced consistently high mortality rates across all sites (median: 84%), reducing mean cover from 5% to 1%. Six years of tracking individual *Pocillopora* colonies revealed strong size-dependent bleaching responses. The heatwave increased the probability of complete and partial mortality more in large than in small *Pocillopora* colonies. Remarkably, elevated partial mortality persisted for at least three years after the thermal stress had subsided, indicating substantial delayed impacts. Contrary to the rapid recovery historically observed on Mo’orea’s reefs following other disturbances, coral cover continued to decline for four years after the bleaching event, with no evidence of recovery. Our findings reveal that bleaching impacts extend far beyond the heatwave itself. Persistent delayed mortality of surviving colonies can limit reef recovery for years, indicating that post-bleaching demographic processes may be a major, yet underappreciated, determinant of coral reefs’ resilience.

## Introduction

Climate change has increased the intensity, frequency, and duration of marine heatwaves (MHW), leading to severe and recurrent coral bleaching and mass mortality events (Hughes et al. 2017; N. J. Holbrook et al. 2019; Oliver et al. 2019). Coral bleaching occurs when the mutualistic symbiosis between the coral host and photosynthetic dinoflagellates (Symbiodiniaceae) breaks down during stressful conditions (Hoegh-Guldberg 1999). Under normal conditions, these microalgal symbionts provide 60 - 70% of the coral’s metabolic energy demand by translocating photosynthetically fixed carbon (Muscatine, McCloskey, and Marian 1981; Grottoli, Rodrigues, and Palardy 2006). If alternative energy sources, such as stored reserves or heterotrophy, are insufficient, prolonged bleaching can lead to starvation and, ultimately, mass mortality (Grottoli, Rodrigues, and Palardy 2006; Anthony et al. 2009). Such mass mortality events can alter species composition and compromise critical ecosystem services for millions of people (Sing Wong, Vrontos, and Taylor 2022). MHW triggered by El Niño-Southern Oscillation have already caused four global bleaching events, resulting in an extensive decline in coral cover worldwide (Claar et al. 2018; Reimer et al. 2024). Given the observed and projected global degradation of coral reefs under ongoing global warming, a better understanding of their immediate and long-term responses to heat stress is crucial.

The intensity of bleaching and mortality generally increases with the level of accumulated heat stress (Sully et al. 2019). Yet, reefs separated by only a few kilometres can experience different thermal regimes due to interactions with currents and local upwelling (McClanahan and Muthiga 2021; Dixon et al. 2022), generating fine-scale variation in heat stress exposure during MHW. Consequently, the bleaching response can differ between adjacent back-reef pools (Thomas et al. 2018) and between neighbouring reefs on a single island (Winslow et al. 2024).

Thermotolerance varies considerably between (Loya et al. 2001; Van Woesik et al. 2011) and within (Van Woesik et al. 2012; Speare et al. 2022; Winslow et al. 2024) coral taxa, resulting in highly variable bleaching responses. Fast-growing branching genera, such as *Acropora* and *Pocillopora*, are generally considered more susceptible to heat stress than slow-growing, massive species, such as *Porites* (Loya et al. 2001; Van Woesik et al. 2011). These differences may partly be due to life-history trade-offs between growth and stress tolerance, and to differences in heterotrophic plasticity (Grottoli, Rodrigues, and Palardy 2006) and energy reserves (Grottoli et al. 2014). Additionally, Symbiodiniaceae taxa differ in heat tolerance, contributing to differences in bleaching susceptibility of the coral holobiont (Rowan 2004; Berkelmans and Van Oppen 2006). The ability to shuffle or switch the composition of Symbiodiniaceae communities varies between different coral species (Putnam et al. 2012; Voolstra and Ziegler 2020), potentially contributing to variations in thermotolerance (García et al. 2024).

The bleaching response also differs greatly within the same coral species (Humanes et al. 2022; Naugle et al. 2024). This variability can result from genotype-specific associations with Symbiodiniaceae of different heat tolerances (Kinzie et al. 2001) and with different bacterial taxa (Voolstra and Ziegler 2020). Differences in colony size are associated with differences in bleaching prevalence, severity, and mortality (Brandt 2009; Burn et al. 2023; Speare et al. 2022; Winslow et al. 2024). Larger corals are often more susceptible to bleaching and mortality than smaller conspecifics, possibly because slower diffusive exchange in larger colonies hinders the removal of harmful metabolites produced during bleaching (Nakamura and Van Woesik 2001; Van Woesik et al. 2012). Conspecific differences in thermotolerance can also result from different thermal stress histories (Glynn et al. 2025) and ecological memory (Hackerott, Martell, and Eirin-Lopez 2021). The underlying mechanisms are still poorly understood (Vompe et al. 2024), but the pre-exposure to variable temperatures (DeMerlis et al. 2022; Brown, Martynek, and Barott 2024; Ferrara et al. 2024) and repeated heat stress (Speare et al. 2022; Nicholas A. J. Graham et al. 2024) can increase coral thermotolerance through acclimatisation (Hackerott, Martell, and Eirin-Lopez 2021; Howells et al. 2013; Van Oppen et al. 2018) and adaptation (Van Oppen et al. 2015). Colony size may therefore act in opposing directions: larger colonies may carry a longer exposure history that confers thermotolerance, but they may also be more prone to metabolic stress. The net effect on survival is unclear. Considering the importance of large, fecund corals (Hall and Hughes 1996) for reef recovery through recruitment (Hughes et al. 2019), it is crucial to better understand how MHW drive size-dependent bleaching responses.

The time between bleaching events has decreased dramatically (from an average of 27 years in the early 1980s to 5.9 years in 2016), and this trend is expected to continue over the next few decades (Hughes et al. 2018), giving reefs less time to recover. After disturbances such as a bleaching event, the recovery begins with fast-growing species (Morais et al. 2024) and diversity increases over time (Grigg 1983). This can lead to boom-and-bust cycles (Morais et al. 2024) in which fast-growing but sensitive species lead to a rapid recovery of coral cover that readily collapses upon the next disturbance due to their lower stress tolerance. The recovery rate depends on the intensity of the experienced stress (Connell, Hughes, and Wallace 1997) and the type of disturbance. For example, the recovery after an outbreak of Crown-of-thorns starfish (COTS, *Acanthaster* sp.) was faster than after bleaching events and tropical storms (N. A. J. Graham, Nash, and Kool 2011). Additionally, local conditions, such as nitrogen concentration (Donovan et al. 2020), herbivorous fish biomass, wave exposure, depth (Robinson, Wilson, and Graham 2019), and spatial competition (Kopecky et al. 2024), impact the recovery of reefs. While bleaching-induced mortality is commonly quantified within months of thermal stress, much less is known about whether surviving colonies continue to experience elevated mortality and how such delayed effects influence reef recovery.

By integrating two long-term monitoring datasets, this study aims to determine how fine-scale variation in heat stress exposure shapes coral bleaching responses and recovery following a severe MHW in Mo’orea, French Polynesia. Circular volcanic islands such as Mo’orea are ideal for such studies, as seawater temperatures can differ among neighbouring reefs due to interactions with ocean currents and wind patterns (McClanahan and Muthiga 2021). Following the near-total loss of coral cover in 2010 due to a COTS outbreak and a cyclone, the reefs of Mo’orea showed a remarkable recovery, exceeding pre-disturbance levels after just four years (Adjeroud et al. 2018). In 2019, the highest level of thermal stress since records began in 1985 (see SI) triggered a bleaching event that caused widespread coral mortality (Winslow et al. 2024; Wyatt et al. 2023). We examined the small-scale variations in accumulated heat stress during this bleaching event and how these differences may have impacted mortality and recovery among coral genera. Based on a study of the same reefs (Winslow et al. 2024), we hypothesised that *Acropora* corals were more affected by heat stress than *Pocillopora* corals. While we expected larger corals to be more severely affected by heat stress (Speare et al. 2022; Winslow et al. 2024), the implications for recovery were unknown. Therefore, we examined how colony size affected coral growth and mortality before, during, and after the bleaching event.

## Materials & Methods

### Study sites

Mo’orea (French Polynesia) is a volcanic island within the Society Archipelago in the South Pacific (17.54° S, 149.83° W, Figure 1). In this study, we focus on the outer reefs, which are characterised by steeply declining slopes and dominated by *Pocillopora*. Two datasets were analysed in this study to *i)* assess island-wide patterns of mortality and recovery after the 2019 bleaching event triggered by a MHW (“Mo’orea” data) and *ii)* examine the impact of colony size on bleaching-induced mortality and growth (“Polynesia Mana” dataset). Island-wide patterns were assessed by examining changes in coral cover at 13 sites, four of which were located on the west coast (Ha’apiti, Taotaha, Tetaiuo, and Gendron), four on the north coast (Tiahura, Entre deux baies, Piha’ena, and Aroa), and five on the east coast (Nuarei, Tema’e, Motu Ahi, Afareaitu, and Ma’atea, Figure 1). The impact of colony size was investigated at two sites, Entre deux baies on the north coast and Ha’apiti on the west coast (Figure 1).

**Figure 1:**
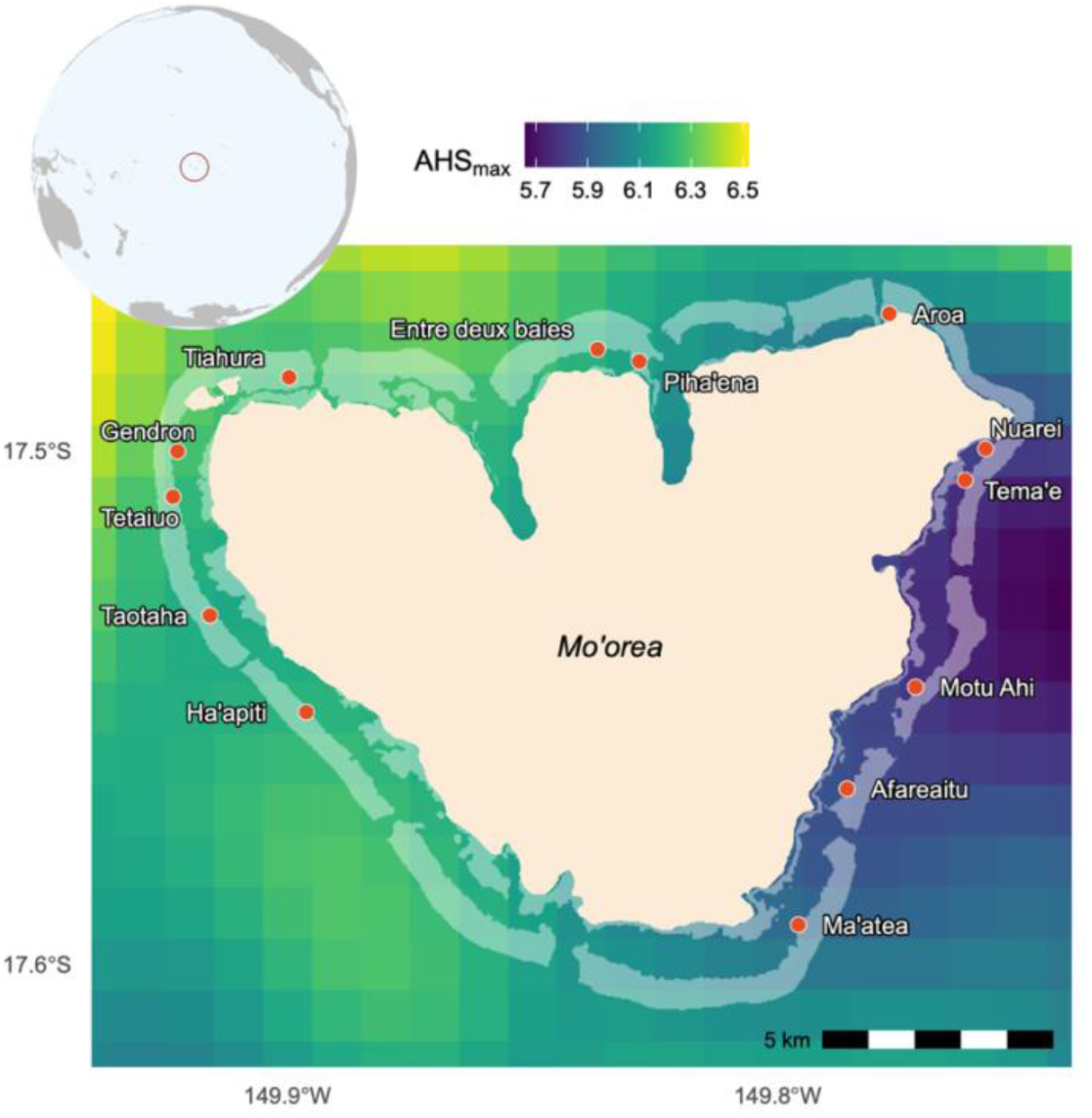
Map of Mo’orea and surveyed sites. Top left: Mo’orea in the Central Pacific. Main plot: The raster layer in the background visualises the maximum accumulated heat stress (AHS_max_ in °C weeks) during the 2019 heatwave.

### Fine-scale patterns in accumulated heat stress

An adaptation of the Degree Heating Weeks approach (Skirving et al. 2020) was used to calculate accumulated heat stress (AHS) throughout the 2019 bleaching event. This metric is based on the Multi-scale Ultra-high Resolution (MUR) Sea Surface Temperature (SST) dataset version 4.1 with a spatial resolution of 1 km (Chin, Vazquez-Cuervo, and Armstrong 2017). In 2019, AHS was calculated as the cumulative sum of SST values above 29.0 °C, the bleaching threshold for corals in Mo’orea (Pratchett et al. 2013), within a moving 84-day window and scaled by 1 week. The maximum value in 2019 (AHS_max_) was extracted for each site.

### Island-wide patterns of coral mortality and recovery

Benthic surveys for the “Mo’orea MPA” time series (Moritz et al. 2021; SNO CORAIL 2025) were conducted annually (Figure 2 and Table SI-1) at a depth of 10 m across 13 sites (Figure 1) using the point-intercept method (Hill and Wilkinson 2004). At each site, the type of benthic cover (for corals at the genus level) was recorded every 50 cm along a 25 m transect (50 points per transect, 3 transects per site). In this study, we focused on surveys carried out between 2019, just before the onset of the bleaching event (Figure 2), and 2023, and on the three most abundant coral genera (mean ± SE in 2019), i.e. *Pocillopora* (35.2% ± 2.7%), *Montipora* (7.2% ± 1%), and *Acropora* (5.0% ± 0.6%). Data from 2024 were not included because a recent outbreak of COTS led to almost complete mortality of the outer reefs in Mo’orea (personal observation).

**Figure 2:**
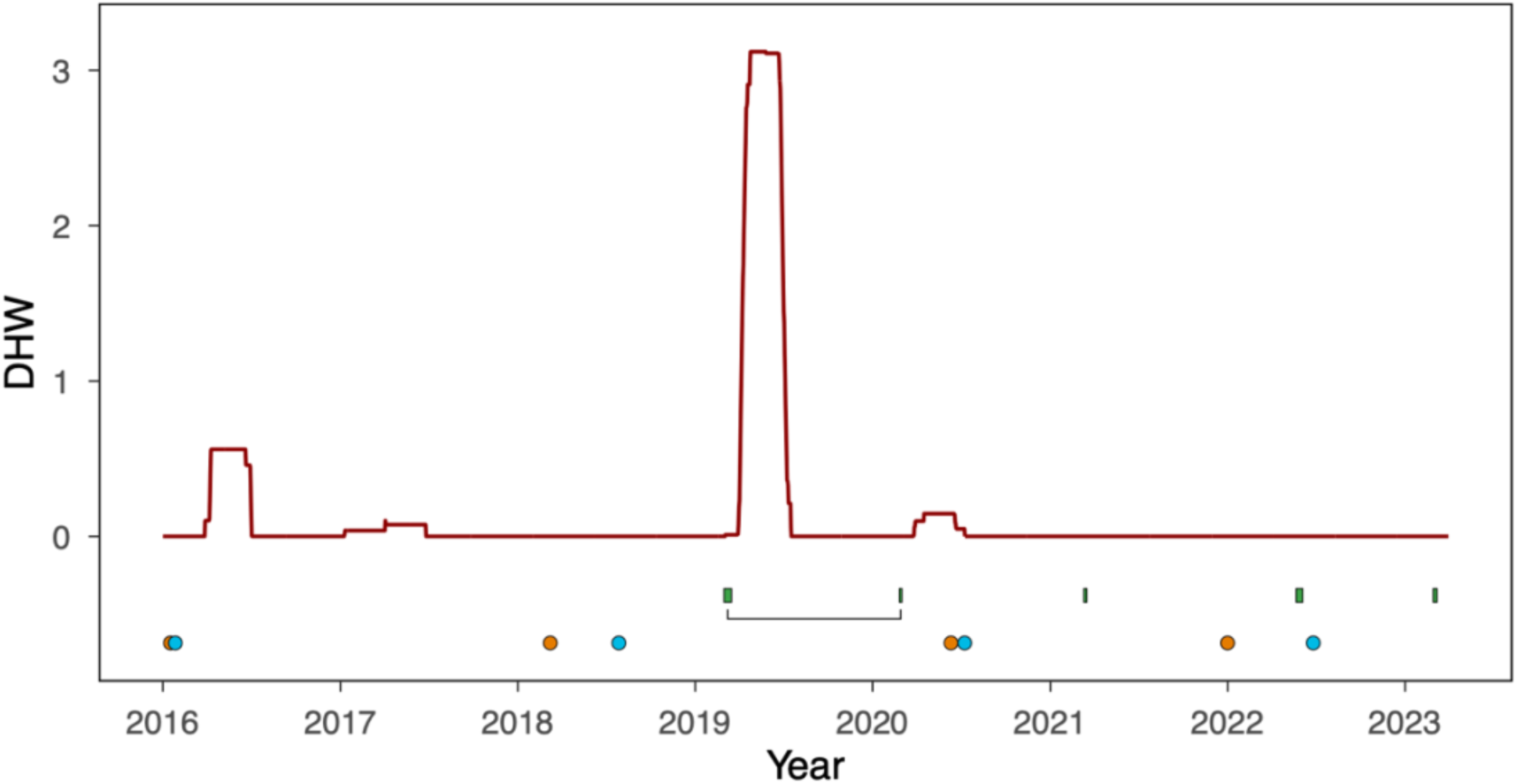
Marine heat stress and sampling times. The red line shows the accumulated heat stress, which is expressed as the mean degree heating weeks around Mo’orea. The data were downloaded directly from the NOAA CRW website (see Supplementary Information for details). The green squares show the sampling periods during which the benthic transects were performed. The bracket shows the comparison used to assess immediate mortality after the bleaching event. The orange and blue points denote the sampling times for the permanent photo quadrat dataset (orange for Ha’apiti and blue for Entre deux baies).

**Figure 3:**
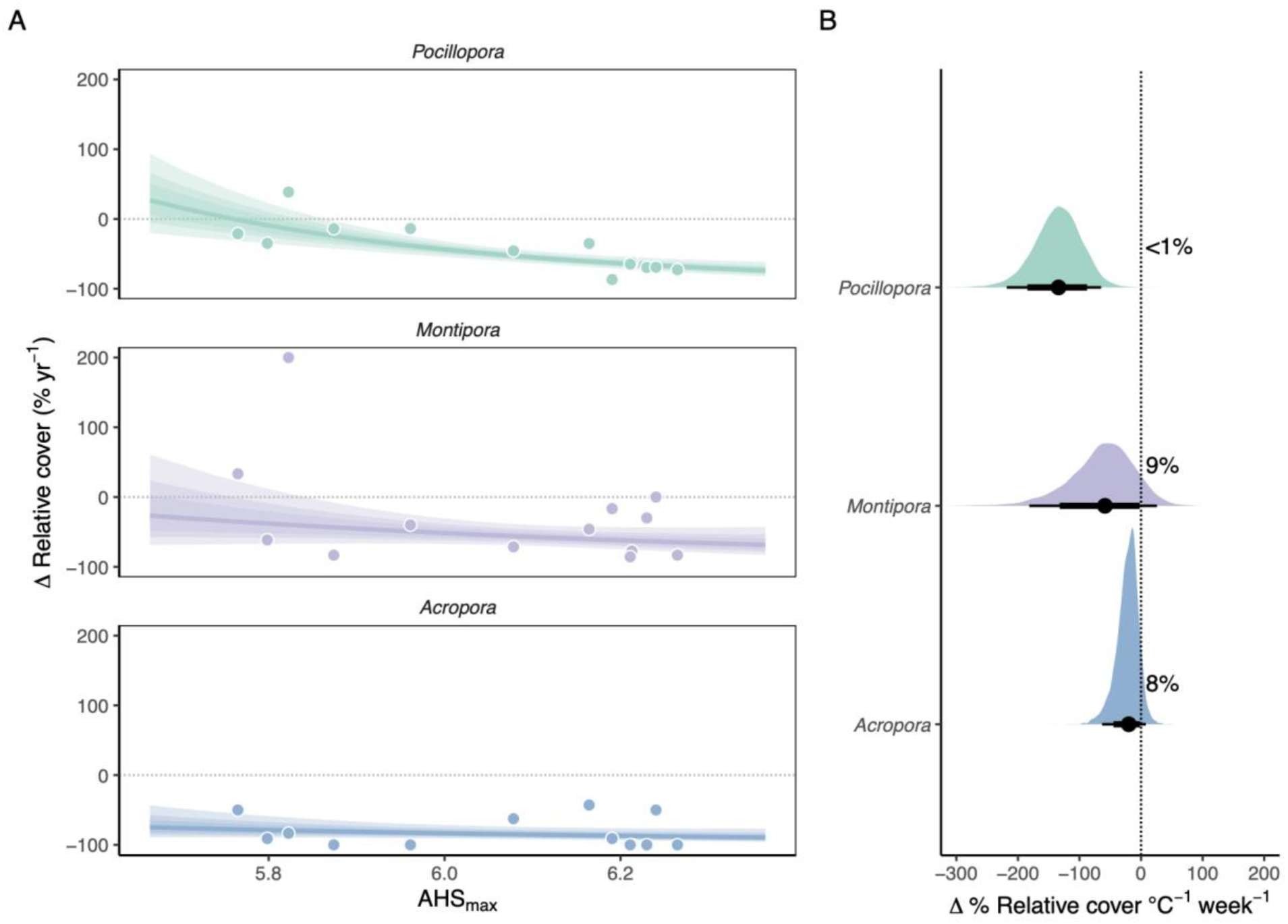
**A:** Model predictions of the relationship between relative change in coral cover between 2018 and 2020 and maximum accumulated heat stress (AHS_max_, in °C weeks) during the 2019 heatwave. The shaded area corresponds to the 60%, 80%, and 95% credible intervals (CI) of the posterior predictive distribution, with the line representing the median. **B:** shows the posterior predictive distribution for the average slope over the range of AHS_max_. The point below the density plot represents the median, and the horizontal lines the 80% and 95% CI. The numbers next to the dashed line show the fraction of the distribution above 0.

The immediate bleaching-induced mortality was assessed with a Bayesian binomial model with a logit-link function (*brms* package, Bürkner, 2017) in R (version 4.4.2, R Core Team, 2024). This model estimated the proportion of the 50 points per transect that were on living corals of different genera directly before (March 2019) and after the bleaching event (March 2020). The interaction of AHS_max_, year (coded as a factor), and the coral genus (*Pocillopora*, *Montipora*, and *Acropora*) was used as explanatory variable. Spatial dependencies were accounted for by including a random slope for each site (*1 + year*|*site*). Broad normal priors with a mean of zero and a standard deviation of ten were used for the slope and intercept. The Hamiltonian Monte Carlo algorithm used four chains with 4,000 iterations after a warm-up period of 2,000 iterations. Model convergence was assessed using R^^^ values (*R*^^^ ≈ 1.0). Additionally, posterior predictive checks (*pp_check()*) were performed, and the conditional R^2^ value (including fixed and random effects, 0.85) and the marginal R^2^ value (only fixed effects, 0.74) were calculated using the *r2_bayes()* function in the *performance* package (Lüdecke et al. 2021). The posterior predictive distribution was calculated with the *epred_draws()* function (Kay 2023) to determine the relative change in coral cover compared to the cover in 2019 (i.e. cover in 2019 = 100%). These posterior predictive distributions were calculated for each genus along the range of AHS_max_ values that corals were exposed to in Mo’orea. Due to the non-linear relationship between the relative change in coral cover and AHS_max_, the posterior predictive distribution for the average slope across the range of AHS_max_ values was calculated for comparisons between coral genera.

To assess to what extent the reefs recovered in the years following the bleaching event, the coral cover was estimated for each year by using the interaction of AHS_max_, genus, and year (coded as a factor for each year between 2019 and 2023) as explanatory variables and site, genus, and the interaction of year, site, and genus as random factors [*(1 | site) + (1 | genus) + (1 | year:site:genus)*]. As described above, the coral cover was estimated as the proportion of the 50 points per transect that were on living corals of the three genera. Weakly informative priors following a normal distribution with a mean of zero and a standard deviation of ten were used for the intercept and slope. The same number of chains and iterations and the same model checks as described for the models above were used. The conditional R^2^ was 0.89, and the marginal R^2^ was 0.57. The posterior predictive distribution for the remaining coral cover relative to 2019 was calculated for each year and for the minimum (5.76 °C weeks, for Tema’e), mean (6.06 °C weeks), and maximum (6.26 °C weeks, for Gendron) value of AHS_max_ (Figure 4), ignoring the random effects (*re_formula = NA*), as described above. Additionally, the relative change in coral cover at each site was calculated based on the AHS_max_ at that site (Figure 5), incorporating random effects (*re_formula = NULL*).

**Figure 4:**
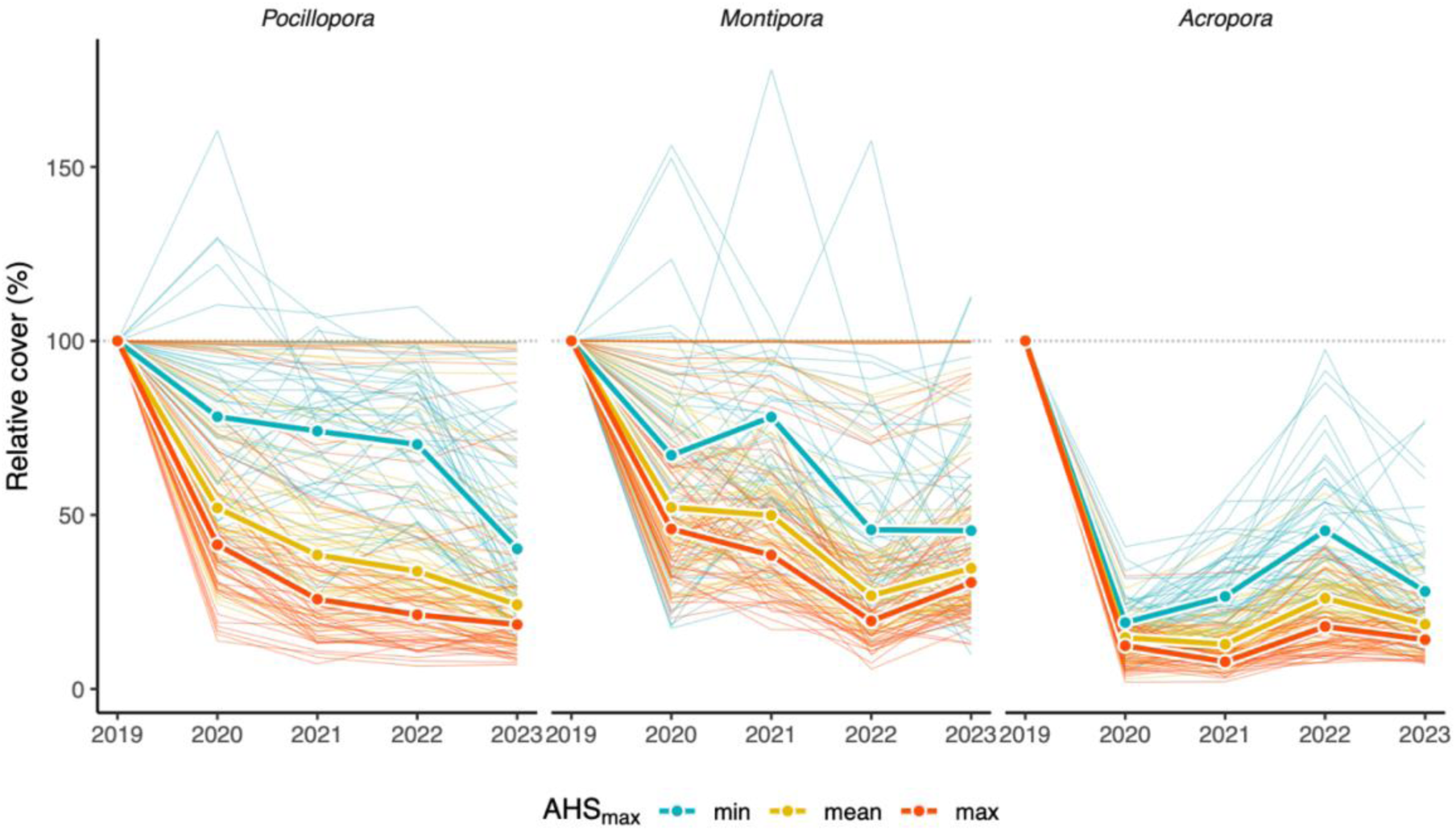
Predicted benthic cover (in percent) relative to the cover in 2019 (dashed line at 100%) for *Pocillopora*, *Montipora*, and *Acropora* at different values for the highest accumulated heat stress in 2019 (AHS_max_). min = minimum AHS_max_ value (5.76 °C weeks, in green), mean = mean AHS_max_ value (6.06 °C weeks, in yellow), max = maximum AHS_max_ value (6.26 °C weeks, in red). The thick lines indicate the median of the posterior predictive distribution, and the thin lines represent 100 randomly chosen draws from the posterior predictive distribution to visualise the variability.

**Figure 5:**
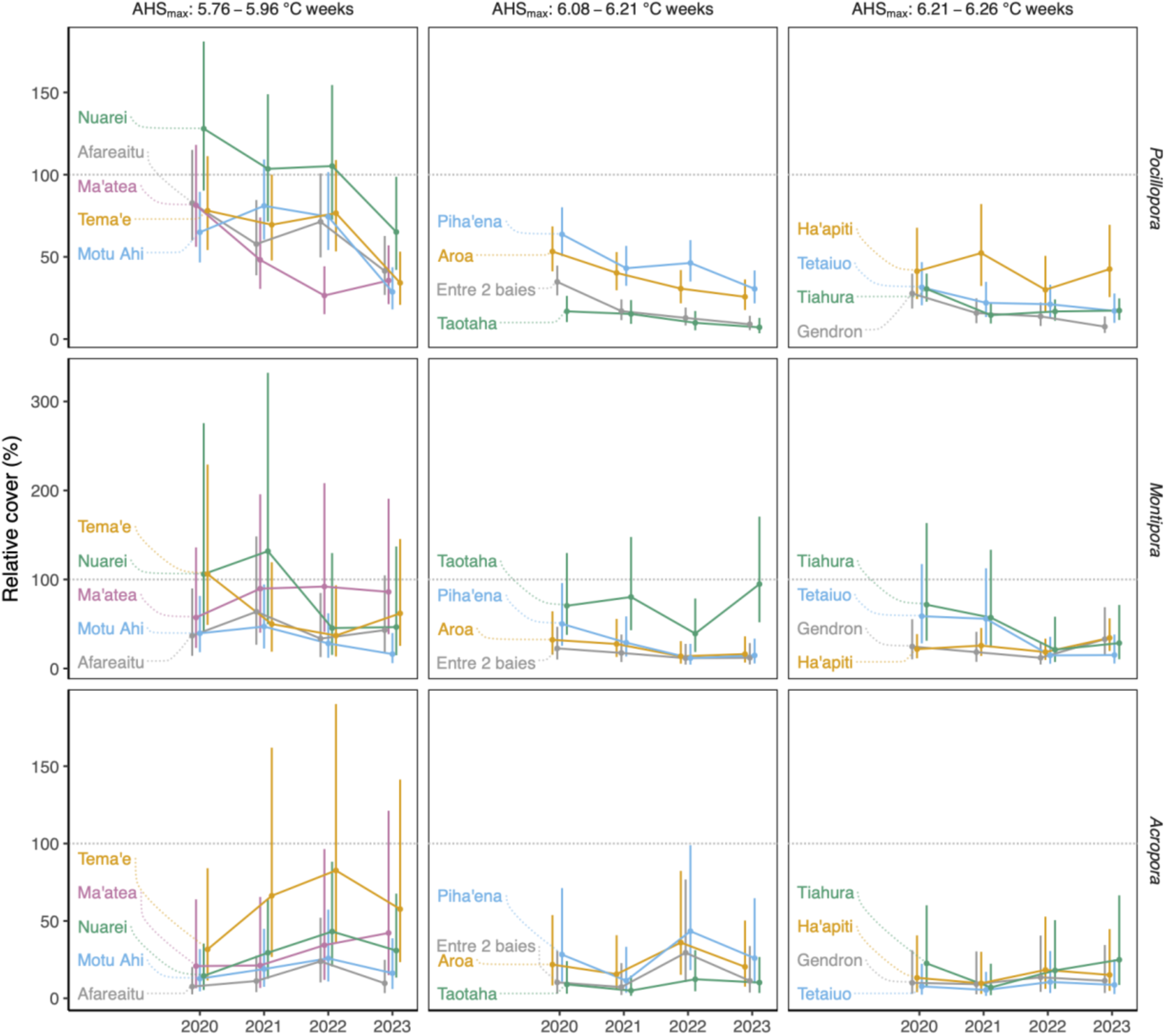
Predicted benthic cover (in percent) relative to the cover in 2019 (dashed line at 100%) for *Pocillopora*, *Montipora*, and *Acropora* at the different sites (rows). The sites were grouped (columns) according to the values for the highest accumulated heat stress in 2019 (AHS_max_, in °C weeks). Within each column, different colours indicate different sites. The points show the median and the vertical lines the 89% credible intervals from the posterior predictive distribution.

### Size-dependent coral mortality and growth

As part of the “Polynesia Mana” long-term monitoring programme (SNO CORAIL 2025), surveys using permanent photo quadrats were conducted every two years from 2016 to 2022 (Figure 2, Table SI-1) to assess the impact of colony size on bleaching mortality and recovery. A rope was stretched at a depth of 10 m between two permanently fixed metal poles 20 m apart. A 1 x 1 m quadrat was photographed at one-meter intervals along the rope (20 quadrats per site), enabling the quadrats to be placed in the same spot for each survey.

Photographs at two study sites (Entre deux baies in the north and Ha’apiti in the west) were analysed to quantify size-dependent mortality and growth during the 2019 bleaching event and the subsequent recovery. Colony size was only estimated for *Pocillopora* corals (19% average benthic cover on photo quadrats between 2016 and 2022 at both sites), given the low cover of other genera (*Montipora*: 2.9%, *Porites*: 1.9%, *Acropora*: 1.7%). These low cover values meant that replication would have been insufficient for the analysis described below. The living surface area of all *Pocillopora* colonies (n = 3090) was manually annotated with *TagLab* (Pavoni et al., 2022, version 2023.5.17) for 2016, 2018, 2020, and 2022. Data from 2024 were not included in this study due to the COTS outbreak mentioned above. Each colony was identified on the corresponding photos of the following surveys and was considered dead if the colony disappeared or no living tissue remained. TagLab’s *compute genets* function was used to give each colony a unique ID and export the area for each year the colony was present. For pairs of years (2016 - 2018, 2018 - 2020, and 2020 - 2022), the probability of growth and mortality of each colony was estimated. If the surface area increased by more than 5%, the colony was considered to have grown; if it decreased by more than 5%, the category “partial mortality” was assigned. The buffer of 10% was assumed to account for inaccuracies during annotation and projection errors when representing three-dimensional coral as a two-dimensional surface. If the area changed by less than 5% in either direction, the “no change” category was assigned. The probability of a colony being assigned to each category was estimated using a Bayesian Generalized Additive Model, with logged surface area as explanatory variable and the categorical family with a logit link-function in *brms*. Different smoothing terms were considered for each site [*s(log_area, by = site)*]. A separate model was used for each comparison (2016 - 2018, 2018 - 2020, and 2020 - 2022). A random intercept for the quadrat ID was added to account for spatial dependencies. The same number of chains and iterations and the same model checks as described for the models above were used. The posterior predictive distributions for both sites were combined. A threshold was calculated for the maximum colony size corresponding to a probability for complete mortality of at least 50% as well as the minimum colony size corresponding to at least 50% partial mortality.

All data and statistical analyses can be accessed on GitHub https://github.com/andieich/bleaching_2019

## Results

### Fine-scale variation in the magnitude of heat stress

The MHW began in March 2019, peaked in May, and declined in July (NOAA CRW degree heating week metric, see SI and Figure 2). The fine-scale spatial patterns of exposure to heat stress were assessed by comparing the maximum accumulated heat stress (AHS_max_). The AHS_max_ differed across Mo’orea and was generally lower in the east than in the north and west. The mean AHS_max_ was 6.06 °C weeks, the minimum 5.76 °C weeks in Tema’e, and the maximum 6.26 °C weeks in Gendron (Figure 1).

### Heat stress-related mortality varies among coral genera and the intensity of heat stress

The direct impact of the heat stress was evaluated by quantifying changes in coral cover one year after the heatwave relative to pre-bleaching levels (Figure 2, Table SI-1). The mean coral cover across all sites declined from 35% to 17% for *Pocillopora*, from 7% to 3% for *Montipora,* and from 5% to 1% for *Acropora*. Since coral cover varied considerably among sites, changes were expressed relative to pre-bleaching values, with values below zero corresponding to mortality and values above zero corresponding to recovery.

The median relative change across all sites (posterior predictive distribution) was −46% (95% CI: −76%, 37%) for *Pocillopora*, −54% (95% CI: −76%, 4%) for *Montipora*, and −84% (95% CI: −93%, −63%) for *Acropora*. The immediate mortality of *Pocillopora* after the bleaching event increased with AHS_max_ (i.e., cover decreased, Figure 3, median slope: −134, 95% CI: [−218, −65] Δ % relative cover °C^-1^ week^-1^). There was strong evidence for this relationship, with the posterior probability that the slope was negative exceeding 99%. For *Montipora*, bleaching-induced mortality varied across the range of AHS_max_, resulting in a broader posterior distribution of the slope (median: −59, 95% CI: [−182, 26] Δ % relative cover °C^-1^ week^-1^) and a lower posterior probability of 91% to be negative. For *Acropora,* the relationship with AHS_max_ was weaker (median slope: −20, 95% CI: [−63, 8] Δ % relative cover °C^-1^ week^-1^) with a posterior probability of 92% that the slope is negative.

### No sign of recovery after mass bleaching mortality

No substantial recovery was observed for the three dominant coral genera on the reefs of Mo’orea within the four years following the bleaching event. First, we predicted the general impact of AHS_max_ on coral cover relative to before the bleaching event, ignoring site-specific variability (Figure 4). Instead of recovery, we observed an overall decline in cover for all three genera, which was stronger at sites exposed to higher levels of heat stress during the 2019 bleaching event. Overall, mortality remained elevated for at least three years after the marine heatwave.

*Pocillopora* cover declined continuously after the bleaching event, and mortality was lower at the minimum level of AHS_max_ than at mean and maximum exposure levels. Although this trend was observed for all years, the loss in coral cover was most severe at the minimum AHS_max_ between 2022 and 2023 (Figure 4, Table SI-2). A similar decline was observed for *Montipora* but with higher variability (Figure 4, Table SI-2). *Acropora* cover collapsed immediately after the bleaching event, regardless of the intensity of the heat stress experienced, and increased slightly in cover from 2020 to 2022. While consistent at multiple sites (Figure 5), the change in absolute cover was minor. Of the 1,950 points across all transects per year combined (13 sites, 3 transects, 50 points per transect), 97 points with *Acropora* were found before the bleaching event (mean cover = 5.0%), compared to 16 in 2020 (0.8%), 17 in 2021 (0.9%), 30 in 2022 (1.5%) and 21 in 2023 (1.1%), with high within-site variability. The highest increase, from 2021 to 2022, was 13 points across all transects (Δ 0.7%). This means that, on average, 0.3 more points with *Acropora* were found per transect (total 50 points) in 2022 than in 2021.

The same model was used to estimate changes in coral cover at the different sites (Figure 5, Table SI-3). At sites with low exposure to accumulated heat stress (AHS_max_ = 5.76 - 5.96 °C weeks), changes in coral cover were more variable for all three genera. At sites with medium (AHS_max_ = 6.08 - 6.21 °C weeks) and high exposure to heat stress (AHS_max_ = 6.21 - 6.26 °C weeks), the coral cover collapsed immediately after the bleaching, although changes in cover were highly variable for *Acropora* and *Montipora*. For example, *Montipora* cover at Taotaha increased from 2022 to 2023, but at very low absolute cover (from an average of 2.3 to 5.0 of the 50 points per transect), challenging the resolution of the transects described above.

### Large colonies are more severely affected by heat stress

The probability of colony growth and mortality before, during, and after the bleaching event was estimated for *Pocillopora* using photo quadrats at two locations, Entre deux baies and Ha’apiti (Figure 1 and SI-2). The number of living *Pocillopora* decreased between 2016 and 2022 by 70% (Figure 6), with the strongest decline (49%) observed between 2020 and 2022. For all years, the size distributions were left-skewed, revealing an overrepresentation of small colonies (Figure 6). From 2016 to 2018, the median colony size (planar area in cm^2^) increased from 38 cm^2^ to 71 cm^2^ and further declined to 51 cm^2^ and 30 cm^2^ for 2020 and 2022, respectively.

**Figure 6:**
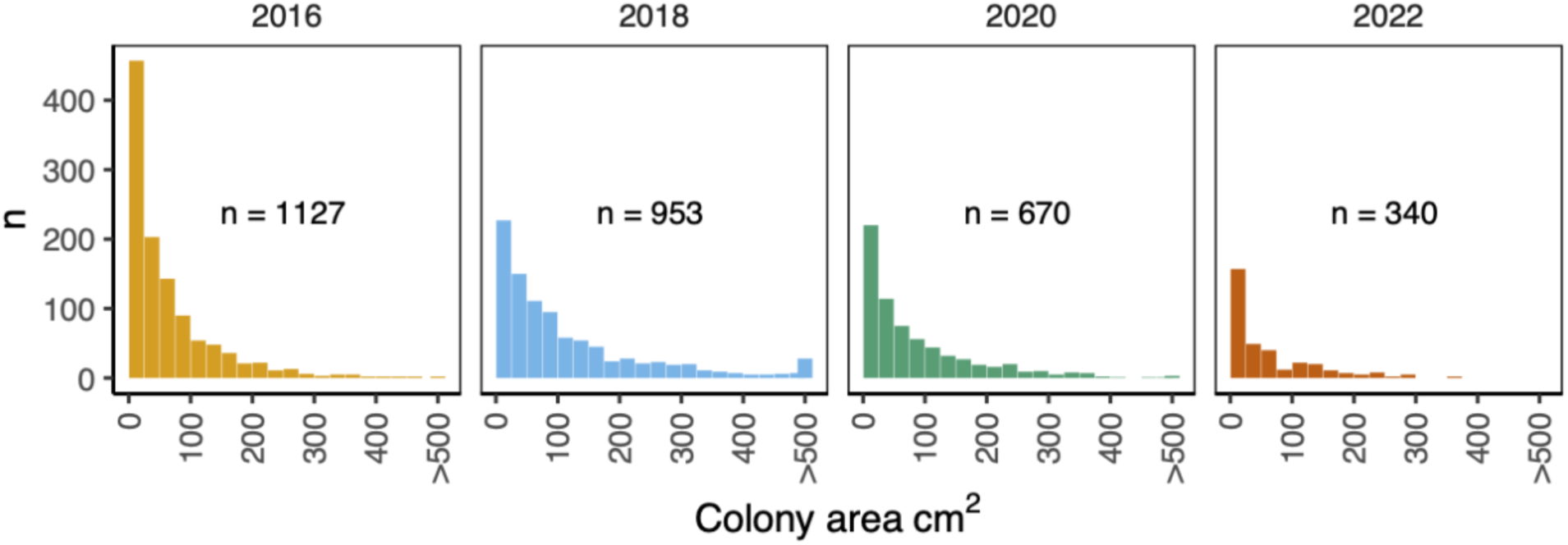
Size distribution for *Pocillopora* corals at Entre deux baies and Ha’apiti between 2016 and 2022. The projected surface area of the colonies was divided into bins of 25 cm^2^. The number in each panel represents the total number of *Pocillopora* colonies observed in that year.

The size-dependent probabilities for growth (increase in area), partial mortality (decrease in area), complete mortality, and no change were modelled to better understand how colony size can influence coral bleaching responses during and after a marine heatwave (Figure 7). From 2016 to 2018, a period without high levels of heat stress in Mo’orea, the probability of complete mortality decreased with colony size (Figure 7A), while the probability of growth increased with colony size. The probability for partial mortality was overall low (below a median of 50%) but slightly increased for larger colonies.

**Figure 7:**
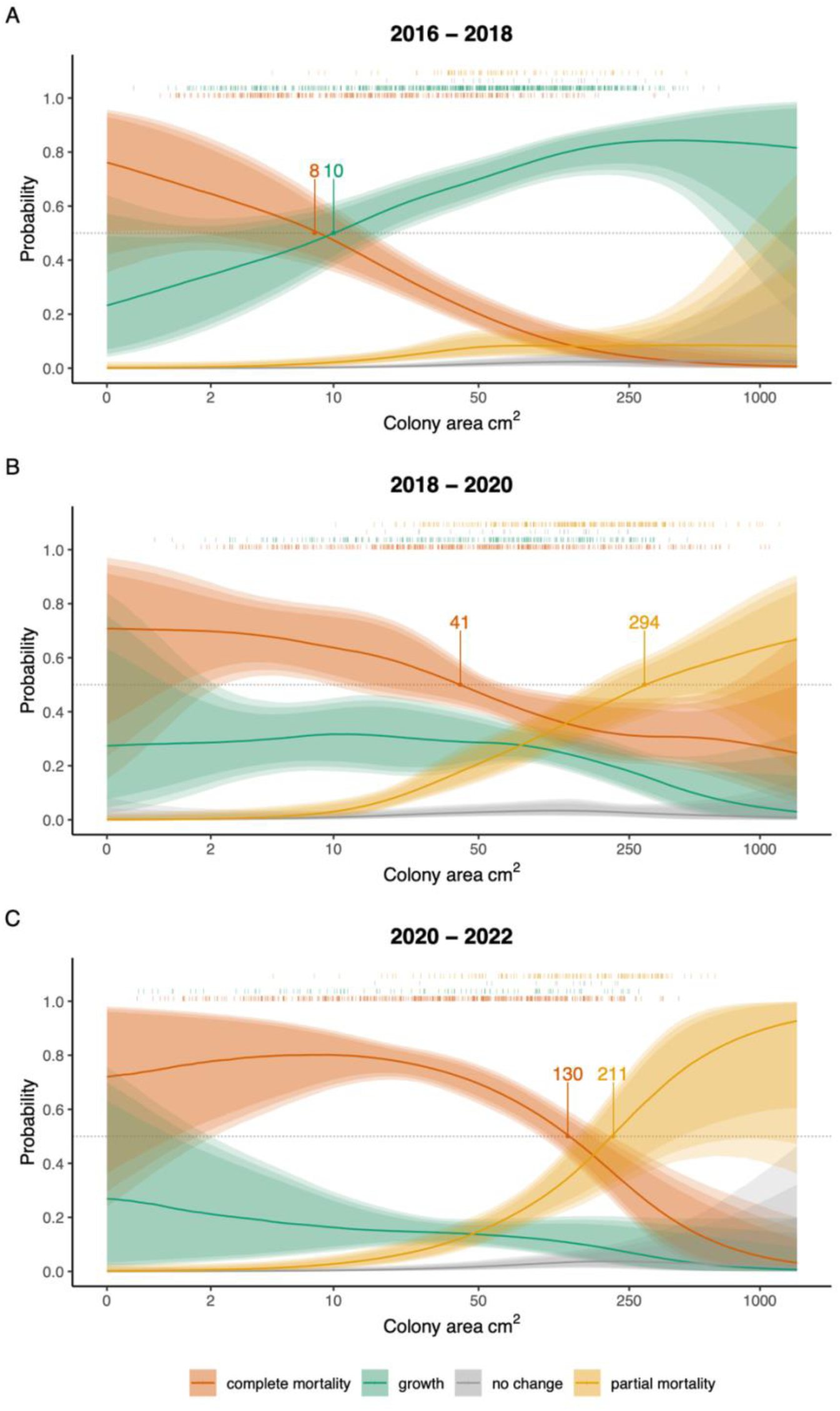
Size-dependent probabilities of growth, complete mortality, partial mortality, and no change for *Pocillopora* at Entre deux baies and Ha’apiti **A** between 2016 - 2018 (before bleaching), **B** 2018 – 2020 (including bleaching), and **C** 2020 – 2022 (after bleaching). The x-axis (log scale) shows the colony area in the first year of the survey. The lines represent the median of the expected values, and the shaded areas represent the 80%, 90%, and 95% credible intervals. The coloured ticks above the plot indicate the occurrences of the respective state category in the raw data. The grey dashed line represents 50% probability, and the annotations represent the corresponding median colony sizes.

The size-dependent impact of the 2019 bleaching event was estimated using a model based on surveys conducted between 2018 and 2020 (Figure 7B). To improve comparability, thresholds were calculated for the maximum and minimum colony sizes corresponding to a probability of complete and partial mortality of at least 50%, respectively, as well as the minimum colony size corresponding to a probability of at least 50% for growth (see annotations in Figure 7). The probability of complete mortality was overall higher compared to years without heat stress and decreased with colony size. However, the decrease in complete mortality was observed for larger corals than before the bleaching event (at least 50% probability of complete mortality for colonies smaller than 41 cm^2^ between 2018 and 2020, compared to 8 cm^2^ between 2016 and 2018, Figure 7). Larger corals were more affected by partial mortality during the heatwave than before (more than 50% probability for partial mortality for colonies larger than 294 cm^2^ between 2018 and 2020, compared to below 50% probability in 2016 and 2018, independent of size, Figure 7). Growth probability was also lower during the bleaching event (below 50%, independent of size, Figure 7) compared to before the bleaching event (more than 50% probability of growth above 10 cm^2^ between 2016 and 2018, Figure 7).

Even after the heat stress had ended, the size-dependent probabilities of growth and mortality between 2020 and 2022 remained similar to the trajectories observed during the bleaching event (Figure 7C). Colonies showing a probability of at least 50% for complete mortality were larger (below 130 cm^2^) than those observed before and during the bleaching event. However, the threshold for 50% probability of partial mortality occurred for smaller colonies (larger than 211 cm^2,^ Figure 7). As during the bleaching event, the probability of growth was generally low, below 50%, and independent of size.

## Discussion

Climate change is leading to more frequent and intense coral bleaching events, with major consequences for reef structure and functioning (Hughes et al. 2017; N. J. Holbrook et al. 2019; Oliver et al. 2019). However, the long-term effects of bleaching on coral populations and reef recovery are not well understood. Therefore, it is important to determine how bleaching-induced declines in coral cover evolve after thermal stress has ended, and how these trajectories vary among reef sites and coral genera. The 2019 marine heatwave, the most intense heat stress since satellite records began in 1985 (see SI), caused widespread coral bleaching and mortality (Speare et al. 2022). By combining island-wide surveys (2019 - 2023) with colony-level tracking of *Pocillopora* (2016 - 2022), we quantified mortality and recovery across spatial (site), temporal (recovery trajectory), and biological (genus and size) scales. Immediate *Pocillopora* mortality increased with heat stress, whereas *Acropora* mortality was uniformly high. An analysis of size-dependent growth and mortality in *Pocillopora*, the dominant coral genus in Mo’orea (Figure SI-1), revealed that large colonies experienced a greater impact of bleaching and persistent stress for up to three years after the heatwave. Contrary to the expectation that reef recovery begins once thermal stress subsides, our results suggest that delayed mortality of surviving colonies can continue to drive declines in coral cover for years, fundamentally altering post-disturbance recovery trajectories.

### Post-bleaching mortality varies among coral genera and sites

Using satellite-derived SST data, we quantified differences in accumulated heat stress around Mo’orea in 2019 at a resolution of 1 km. This allowed us to investigate the impact of maximum accumulated heat stress (AHS_max_) on coral mortality. The lower AHS_max_ along the eastern shore (Figure 1) may be due to stronger currents in the channel between Mo’orea and Tahiti, compared to the northern and western coasts (James et al. 2020).

Consistent with the findings of Winslow *et al*. (2024), *Acropora* experienced widespread and severe mortality during the 2019 bleaching event, with a median mortality rate of 84% across all sites, irrespective of heat stress intensity. In contrast, *Pocillopora* experienced a lower mortality rate (median 46%), which increased with AHS_max_, consistent with a greater thermotolerance than *Acropora*. These contrasting responses may reflect differences in microbiome flexibility. *Pocillopora* is considered a microbiome regulator that maintains relatively stable microbial communities, whereas *Acropora* is a microbiome conformer whose microbial assemblages shift in response to environmental conditions (Ziegler et al. 2019). Such shifts may promote dysbiosis (Boilard et al. 2020), characterised by a transition from mutualistic to pathogenic microorganisms under stress (Maher et al. 2020). Ultimately, this increases the risk of mortality. Similar mechanisms may occur in algal symbiont communities, where the higher Symbiodiniaceae diversity in *Acropora* could increase intra-community competition and reduce host performance during heat stress (Huffmyer et al. 2025).

The weak relationship between heat stress and *Acropora* mortality may have resulted from the low cover of this genus, which increased variability among sites. Similarly, *Montipora* cover declined markedly after bleaching, despite its presumed high thermotolerance, which has been linked to higher energy reserves (Grottoli, Rodrigues, and Palardy 2006). However, its low cover at most sites prevented a robust assessment of its relationship with heat stress intensity and subsequent recovery.

### Large *Pocillopora* colonies suffered greater mortality

Between 2016 and 2022, permanent photo quadrats were used to track the growth and probability of mortality of individual *Pocillopora* colonies at two sites. In years without significant heat stress, mortality decreased with colony size, consistent with previous studies (Connell, 1973). High mortality among small colonies likely reflects a trade-off between growth and stress tolerance (Edmunds, Cheh, and Burgess 2025) and is characteristic of the demographic strategy of many r-selected corals (Vermeij and Sandin 2008). However, partial mortality increased with colony size, probably due to an increased susceptibility to mechanical damage (Madin et al. 2014). Growth probability also increased with colony size, although this pattern may partly be due to the greater detectability of size changes in larger colonies in the photo quadrats.

The 2019 bleaching event mainly affected larger *Pocillopora* colonies, particularly through partial mortality. This suggests that many large colonies experienced severe stress and bleaching during the heatwave but survived until the post-bleaching survey (one year after the MHW). Greater susceptibility of large corals to thermal stress has been widely reported (Brandt 2009; Burn et al. 2023; Speare et al. 2022; Winslow et al. 2024) and is often attributed to reduced mass transfer efficiency, which limits the diffusion of harmful metabolites during bleaching (Van Woesik et al. 2012; Speare et al. 2022). These constraints may outweigh any potential advantages associated with higher pre-exposure to variable temperatures in older colonies. Furthermore, as the majority of corals alive in 2019 recruited after the near-complete collapse of the outer reef in 2010 (Figure SI-1), the largest colonies may have been dominated by a single *Pocillopora* haplotype (mtORF haplotype 11), which suffered higher bleaching mortality and may have grown faster than other *Pocillopora* haplotypes (Burgess et al. 2021). Variation in thermotolerance among *Pocillopora* haplotypes may therefore have contributed to the observed size-dependent patterns of bleaching mortality. However, the effects of size on thermotolerance can vary among regions and species. For example, in Hawai’i, smaller *Pocillopora meandrina* were more susceptible to bleaching than larger conspecifics (Johnston et al. 2020), whereas in Palau, large corals were more likely to bleach but less likely to die, allowing populations to retain highly fecund adults and recover more quickly (Sommer et al. 2024). In contrast, Mo’orea experienced substantial losses of large colonies, combined with a near-complete mortality of coral recruits during the 2019 bleaching event (Speare et al. 2022).

### Delayed mortality as a hidden phase of bleaching disturbance

Three years after the 2019 bleaching event, large *Pocillopora* colonies continued to experience elevated partial mortality, even though no further major heat stress was recorded (see SI). This suggests persistent sublethal impacts and a progressive loss of living tissue. The probability of complete mortality for large corals also remained higher than pre-bleaching levels. Previous studies have reported delayed post-bleaching mortality, with peak mortality occurring nine to ten months after bleaching in Caribbean reefs (Oxenford, Roach, and Brathwaite 2008), and sublethal effects, namely reduced energy reserves and fecundity, persisting for five months in *Acropora* corals after the 2019 bleaching event in Mo’orea (Leinbach et al. 2021). Additional mortality may have resulted from the physical damage caused by fragments of dead colonies being moved by waves. However, to our knowledge, the high levels of partial and complete delayed mortality of large corals observed up to three years after a bleaching event have not previously been documented.

### Lack of recovery after the bleaching event

Unlike the boom-and-bust recovery patterns reported elsewhere (e.g., Morais et al., 2024), Mo’orea’s outer reefs showed no signs of recovery following the bleaching event. Instead, coral cover declined steadily for several years, particularly at sites exposed to the highest heat stress in 2019 (Figures 4 and 5). This contrasts sharply with the rapid recovery observed after the 2010 COTS outbreak and cyclone at the same reefs (Figure SI-1), when coral cover exceeded pre-disturbance levels within four years (Adjeroud et al. 2018; S. J. Holbrook et al. 2018). The prolonged decline may reflect the combined effects of persistent post-bleaching impact and reduced recruitment, as recovery after COTS outbreaks is generally faster than after bleaching events or tropical storms (N. A. J. Graham, Nash, and Kool 2011). The stronger decline at sites exposed to higher heat stress intensity (Figures 4 and 5) suggests that the absence of recovery is at least partly linked to the 2019 bleaching event. In Mo’orea, recovery following the 2010 disturbances may have been facilitated by cyclone-driven removal of dead coral skeletons, particularly on more exposed northern reefs (S. J. Holbrook et al. 2018). This differs from a study reporting faster recovery when the structure of the reef remained intact (Colgan 1987). In Mo’orea, dead skeletons remaining after the 2019 bleaching event reduced herbivory and promoted the growth of turf and unpalatable macroalgae (Kopecky *et al*., 2024). Turf algae can inhibit coral recruitment (Birrell, McCook, and Willis 2005) and macroalgae are strong competitors (e.g., Johns *et al*., 2018), which can negatively affect coral survival, e.g. through allelopathic substances (Rasher and Hay 2010). Consistent with this, macroalgal cover increased across all study sites (Figure SI-1), and post-bleaching coral recruits suffered high mortality, likely due to competition with macroalgae (Kopecky *et al*., 2024).

Water quality may also influence bleaching responses. Nitrogen concentrations vary around Mo’orea and can exacerbate bleaching, particularly under moderate heat stress (Donovan et al. 2020), potentially contributing to some of the observed variability in coral mortality. However, these effects were documented primarily within the lagoon, where terrestrial nutrient input is stronger than on the outer reefs studied here.

Heat stress events other than the 2019 heatwave are unlikely to explain the observed patterns. Although satellite data indicated a minor heatwave in 2016 in Mo’orea (Figure 2), internal wave cooling substantially reduced seawater temperatures and bleaching severity (Wyatt et al. 2023). Likewise, despite evidence of elevated temperature in 2020 from in situ logger data (Vompe et al. 2024), NOAA Degree Heating Weeks, which is a good predictor for coral bleaching risk (Spady *et al*., 2025), showed no significant increase in heat stress around Mo’orea and bleaching was minimal, affecting only 3% of tracked *Pocillopora* colonies and no *Acropora* colonies (Vompe et al. 2024). Taken together, the temperature data and bleaching observations indicate that the 2019 MHW was by far the most severe bleaching event affecting Mo’orea in recent years (Speare *et al*., 2022; Winslow *et al*., 2024). Overall, the high mortality of coral recruits (Speare et al. 2022), the loss of large fecund colonies and resulting shift toward populations of smaller colonies (Hughes et al. 2019), reduced reproductive output of survivors (Leinbach et al. 2021), and increasing competition with algae (Kopecky et al. 2024) may have undermined reef resilience and contributed to the prolonged absence of recovery following the 2019 bleaching event. Beyond reducing greenhouse gas emissions, management could focus on enhancing local conditions that support coral recruitment and limit macroalgal growth.

## Acknowledgments

This study was funded by the France-Canada Research Fund. AE was supported by graduate funding from the French Ministry of Higher Education and Research (MESR). Long-term monitoring was financially supported by CNRS-SNO CORAIL.

## Author contributions

**Andreas Eich:** Conceptualisation; formal analysis; methodology; software; validation; visualisation; writing – original draft; writing – review and editing. **Yannick Chancerelle**: Data curation; methodology; validation; writing – review and editing. **Valeriano Parravicini**: Formal analysis (supporting); validation; writing – review and editing. **Nicolas Derome**: Writing – review and editing; funding acquisition. **Laetitia Hédouin**: Conceptualisation; writing – review and editing; supervision. **Caroline E Dubé**: Conceptualisation; writing – review and editing; supervision; funding acquisition; project administration.

## Supporting information

### Degree heating weeks

To assess the temporal patterns of heat stress in Moorea, degree heating week (DHW) values were downloaded from the NOAA Coral Reef Watch website (https://pae-paha.pacioos.hawaii.edu/erddap/griddap/dhw_5km.html) for the area shown in Figure 2. The mean value across that area was calculated for each day between 1985 and 2023. This parameter was used to describe the temporal patterns of the bleaching event due to its well-established correlation with bleaching risk (e.g. Spady *et al*., 2025). However, the spatial resolution (5 km) was too coarse to enable fine-scale comparison of accumulated heat stress between sites in Mo’orea. To make a relative comparison, the above approach was used to calculate accumulated heat stress (AHS) based on sea surface temperature (SST) data with a resolution of 1 km. As the SST values and threshold differ between the AHS and DHW methods, the absolute values differ. Nevertheless, we believe that this approach combines the strengths of both data sets, particularly given that AHS values are only employed for relative comparisons within a short timeframe and a specific area, without claiming a general correlation between absolute AHS values and bleaching risk.

---

**Table SI-1:** Dates of surveys for benthic transects (Mo’orea MPA data set, across 13 sites) and photo quadrats (Polynesia Mana data set) in Ha’apiti and Entre deux baies for the years considered in this study.

| Year | Benthic transects | Photo quadrats |  |
| --- | --- | --- | --- |
|  |  | Ha'apiti | Entre deux baies |
| 2016 |  | 09.02. | 11.02. |
| 2018 |  | 21.03. | 09.08. |
| 2019 | 12.03. - 29.03. |  |  |
| 2020 | 04.03. - 11.03. | 18.06. | 16.07. |
| 2021 | 16.03. - 24.03. |  |  |
| 2022 | 24.05. - 08.06. | 04.01. | 28.06. |
| 2023 | 28.02. - 10.03. |  |  |

**Table SI-2:** Predicted benthic cover (in percent) relative to the cover in 2019 for *Pocillopora*, *Montipora*, and *Acropora* for the highest accumulated heat stress in 2019 (AHS_max_) for 2020 to 2023. min = minimum AHS_max_ value (5.76 °C weeks), mean = mean AHS_max_ value (6.06 °C weeks), max = maximum AHS_max_ value (6.26 °C weeks). The numbers represent the median of the posterior predictive distribution and the 95% credible intervals (in square brackets).

| **AHS** | ***Acropora*** | ***Montipora*** | ***Pocillopora*** |
| --- | --- | --- | --- |
| **2020** |  |  |  |
| min | 19 [8, 59] | 67 [26, 132] | 79 [36, 120] |
| mean | 15 [7, 39] | 52 [27, 100] | 52 [28, 100] |
| max | 12 [5, 37] | 45 [20, 100] | 42 [18, 100] |
| **2021** |  |  |  |
| min | 27 [11, 71] | 77 [31, 152] | 74 [34, 114] |
| mean | 13 [6, 38] | 49 [27, 100] | 39 [20, 100] |
| max | 8 [3, 26] | 38 [17, 100] | 26 [11, 100] |
| **2022** |  |  |  |
| min | 46 [20, 97] | 46 [17, 100] | 70 [31, 108] |
| mean | 26 [13, 61] | 27 [13, 100] | 34 [17, 100] |
| max | 18 [8, 48] | 19 [8, 100] | 22 [9, 100] |
| **2023** |  |  |  |
| min | 28 [12, 73] | 45 [17, 100] | 40 [16, 100] |
| mean | 19 [9, 49] | 34 [18, 100] | 24 [12, 100] |
| max | 14 [6, 41] | 30 [13, 100] | 19 [7, 100] |

**Table SI-3:** Predicted benthic cover (in percent) relative to the cover in 2019 for *Pocillopora*, *Montipora*, and *Acropora* at different sites from 2020 to 2023. The numbers represent the median of the posterior predictive distribution and the 95% credible intervals (in squared brackets).

| site | 2020 |  |  | 2021 |  |  | 2022 |  |  | 2023 |  |  |
| --- | --- | --- | --- | --- | --- | --- | --- | --- | --- | --- | --- | --- |
|  | *Acropora* | *Montipora* | *Pocillopora* | *Acropora* | *Montipora* | *Pocillopora* | *Acropora* | *Montipora* | *Pocillopora* | *Acropora* | *Montipora* | *Pocillopora* |
| Afareaitu | 8 [2, 25] | 37 [11, 114] | 83 [56, 123] | 11 [3, 34] | 64 [22, 180] | 58 [36, 92] | 24 [8, 62] | 34 [10, 105] | 71 [46, 109] | 10 [3, 31] | 43 [14, 126] | 41 [24, 69] |
| Aroa | 22 [7, 66] | 32 [13, 75] | 53 [39, 73] | 16 [4, 49] | 28 [11, 65] | 40 [27, 56] | 36 [12, 96] | 14 [4, 37] | 31 [20, 45] | 20 [6, 62] | 16 [5, 43] | 26 [16, 39] |
| Entre 2 baies | 10 [2, 39] | 23 [8, 54] | 35 [25, 47] | 7 [2, 29] | 17 [6, 45] | 17 [10, 26] | 29 [8, 96] | 12 [4, 33] | 13 [7, 21] | 11 [3, 42] | 12 [4, 34] | 9 [5, 16] |
| Gendron | 10 [2, 41] | 25 [8, 66] | 28 [17, 43] | 9 [2, 39] | 18 [6, 49] | 16 [8, 27] | 14 [3, 52] | 12 [3, 36] | 14 [7, 25] | 11 [3, 44] | 33 [12, 82] | 8 [3, 16] |
| Ha'apiti | 13 [3, 53] | 22 [10, 43] | 41 [21, 74] | 10 [2, 39] | 26 [12, 50] | 52 [29, 91] | 18 [5, 65] | 18 [8, 38] | 30 [15, 57] | 15 [4, 57] | 34 [17, 63] | 43 [23, 78] |
| Ma'atea | 21 [5, 81] | 57 [19, 164] | 81 [51, 127] | 21 [5, 84] | 90 [34, 241] | 48 [27, 81] | 34 [9, 122] | 92 [34, 252] | 26 [13, 48] | 42 [12, 156] | 86 [33, 229] | 36 [19, 64] |
| Motu Ahi | 13 [3, 39] | 40 [15, 96] | 65 [43, 97] | 19 [6, 54] | 47 [19, 109] | 81 [56, 117] | 26 [9, 69] | 28 [10, 74] | 74 [51, 109] | 16 [5, 47] | 16 [5, 49] | 29 [16, 48] |
| Nuarei | 15 [4, 43] | 106 [32, 343] | 128 [84, 196] | 29 [11, 76] | 132 [41, 416] | 104 [66, 162] | 43 [17, 102] | 45 [11, 166] | 105 [66, 167] | 31 [11, 80] | 47 [11, 181] | 65 [38, 109] |
| Piha'ena | 28 [9, 87] | 50 [22, 112] | 64 [48, 84] | 11 [3, 41] | 29 [11, 69] | 43 [30, 60] | 43 [15, 119] | 12 [3, 34] | 46 [33, 64] | 26 [8, 81] | 15 [4, 41] | 31 [20, 45] |
| Taotaha | 9 [2, 30] | 70 [33, 149] | 17 [9, 29] | 5 [1, 19] | 80 [37, 169] | 15 [8, 26] | 12 [4, 38] | 39 [16, 94] | 10 [5, 19] | 10 [3, 33] | 95 [45, 197] | 7 [3, 14] |
| Tema'e | 32 [9, 107] | 106 [40, 276] | 78 [50, 120] | 66 [22, 195] | 50 [15, 146] | 70 [44, 108] | 83 [29, 233] | 37 [11, 115] | 77 [49, 118] | 58 [19, 174] | 62 [21, 178] | 34 [19, 59] |
| Tetaiuo | 8 [2, 29] | 59 [24, 139] | 31 [18, 51] | 5 [1, 22] | 56 [23, 128] | 22 [12, 38] | 11 [3, 38] | 15 [4, 44] | 21 [11, 37] | 9 [2, 32] | 15 [4, 47] | 17 [9, 31] |
| Tiahura | 23 [6, 72] | 72 [26, 194] | 30 [21, 42] | 7 [2, 29] | 57 [19, 160] | 15 [9, 23] | 18 [5, 63] | 21 [6, 72] | 17 [10, 26] | 25 [7, 83] | 29 [8, 90] | 17 [11, 27] |
| **Mean** | 14 [3, 66] | 47 [12, 190] | 52 [15, 142] | 12 [2, 90] | 45 [10, 198] | 41 [10, 118] | 26 [5, 120] | 22 [5, 130] | 30 [8, 120] | 18 [3, 97] | 30 [6, 153] | 26 [5, 76] |

**Figure SI-1:**
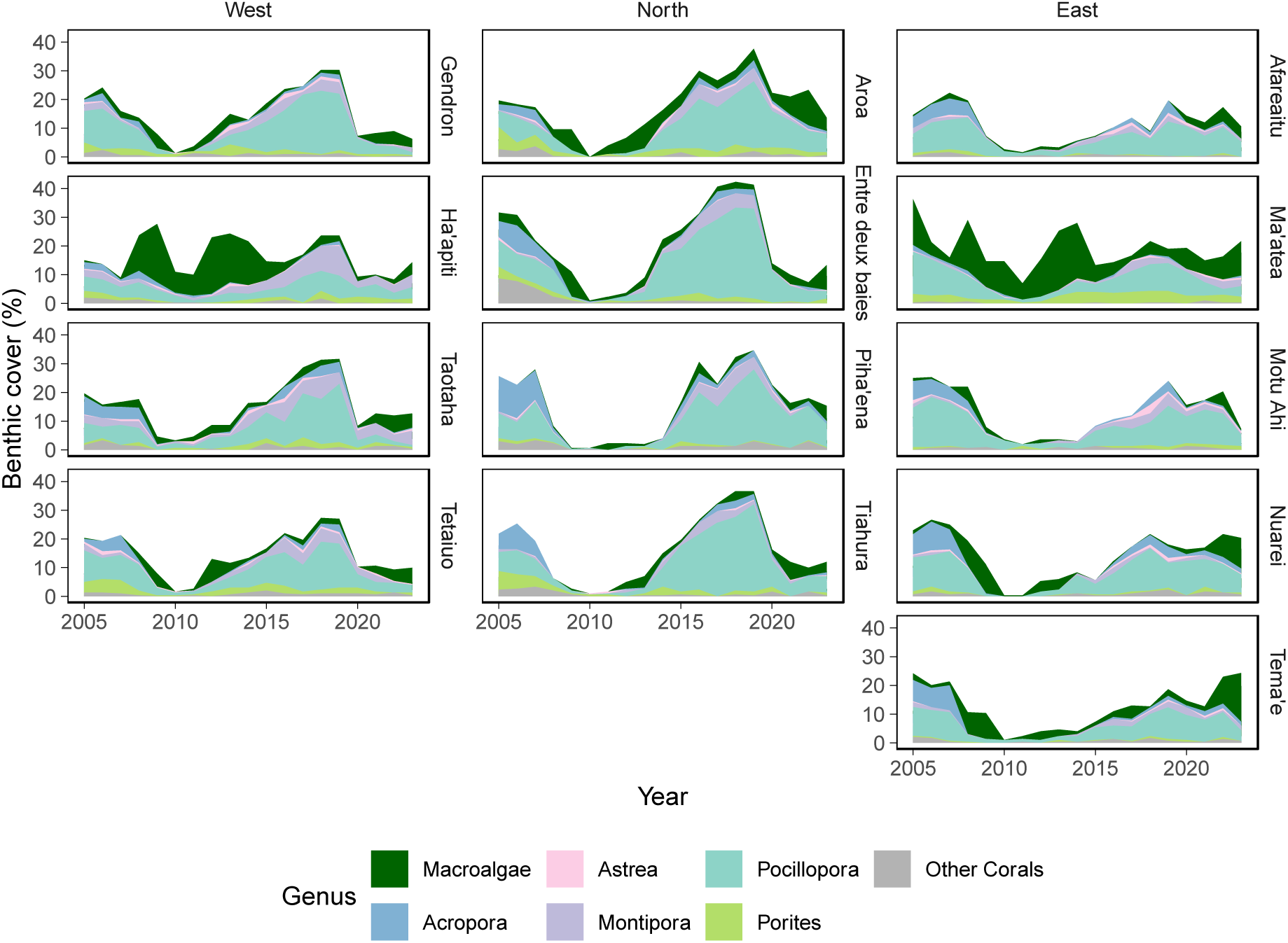
Mean cover of the five most abundant coral genera and for macroalgae between 2005 and 2023.

**Figure SI-2:**
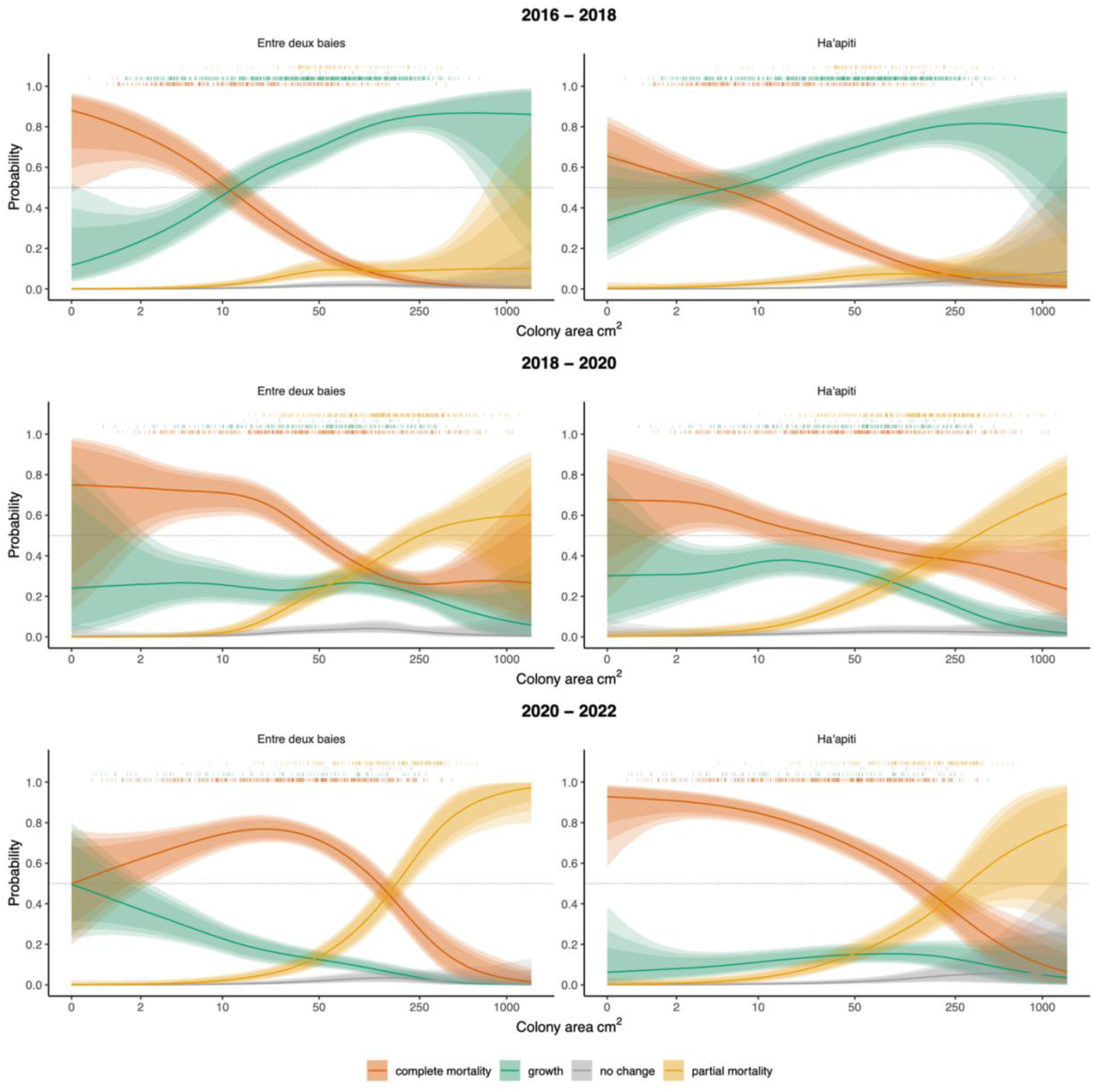
Size-dependent probability of growth, complete and partial mortality, and no change for *Pocillopora* at Entre deux baies (left column) and Ha’apiti (right column) between 2016 - 2018 (top row, before bleaching), 2018 – 2020 (middle row, including bleaching), and 2020 – 2022 (lowest row, after bleaching). The y-axis (on the log scale) represents the colony area in the first year of the survey. The lines represent the median of the expected values, and the shaded area represents the 80%, 90%, and 95 % credible intervals. The coloured ticks above the plot represent the occurrences of the respective state category in the raw data. The grey dashed line represents the 50% probability

## Notes

### Competing Interest Statement

The authors have declared no competing interest.

### Summary of Updates

Correction of minor mistakes and general improvements of writing.

## References

Adjeroud, Mehdi, Mohsen Kayal, Claudie Iborra-Cantonnet, Julie Vercelloni, Pauline Bosserelle, Vetea Liao, Yannick Chancerelle, Joachim Claudet, and Lucie Penin. 2018. ‘Recovery of Coral Assemblages despite Acute and Recurrent Disturbances on a South Central Pacific Reef’. Scientific Reports 8 (1): 9680. 10.1038/s41598-018-27891-3.

Anthony, Kenneth R. N., Mia O. Hoogenboom, Jeffrey A. Maynard, Andréa G. Grottoli, and Rachael Middlebrook. 2009. ‘Energetics Approach to Predicting Mortality Risk from Environmental Stress: A Case Study of Coral Bleaching’. Functional Ecology 23 (3): 539–50. 10.1111/j.1365-2435.2008.01531.x.

Berkelmans, Ray, and Madeleine J. H. Van Oppen. 2006. ‘The Role of Zooxanthellae in the Thermal Tolerance of Corals: A “Nugget of Hope” for Coral Reefs in an Era of Climate Change’. Proceedings of the Royal Society B: Biological Sciences 273 (1599): 2305–12. 10.1098/rspb.2006.3567.

Birrell, Chico L., Laurence J. McCook, and Bette L. Willis. 2005. ‘Effects of Algal Turfs and Sediment on Coral Settlement’. Marine Pollution Bulletin 51 (1–4): 408–14. 10.1016/j.marpolbul.2004.10.022.

Boilard, Aurélie, Caroline E. Dubé, Cécile Gruet, Alexandre Mercière, Alejandra Hernandez-Agreda, and Nicolas Derome. 2020. ‘Defining Coral Bleaching as a Microbial Dysbiosis within the Coral Holobiont’. Microorganisms 8 (11): 1682. 10.3390/microorganisms8111682.

Brandt, M. E. 2009. ‘The Effect of Species and Colony Size on the Bleaching Response of Reef-Building Corals in the Florida Keys during the 2005 Mass Bleaching Event’. Coral Reefs 28 (4): 911–24. 10.1007/s00338-009-0548-y.

Brown, Kristen T., Marcelina P. Martynek, and Katie L. Barott. 2024. ‘Local Habitat Heterogeneity Rivals Regional Differences in Coral Thermal Tolerance’. Coral Reefs 43 (3): 571–85. 10.1007/s00338-024-02484-x.

Burgess, Scott C., Erika C. Johnston, Alex S. J. Wyatt, James J. Leichter, and Peter J. Edmunds. 2021. ‘Response Diversity in Corals: Hidden Differences in Bleaching Mortality among Cryptic *Pocillopora* Species’. Ecology 102 (6). 10.1002/ecy.3324.

Bürkner, Paul-Christian. 2017. ‘brms: An R Package for Bayesian Multilevel Models Using Stan’. Journal of Statistical Software 80 (1): 1–28. 10.18637/jss.v080.i01.

Burn, D., A. S. Hoey, S. Matthews, H. B. Harrison, and M. S. Pratchett. 2023. ‘Differential Bleaching Susceptibility among Coral Taxa and Colony Sizes, Relative to Bleaching Severity across Australia’s Great Barrier Reef and Coral Sea Marine Parks’. Marine Pollution Bulletin 191 (June 2023): 114907. 10.1016/j.marpolbul.2023.114907.

Chin, Toshio Michael, Jorge Vazquez-Cuervo, and Edward M. Armstrong. 2017. ‘A Multi-Scale High-Resolution Analysis of Global Sea Surface Temperature’. Remote Sensing of Environment 200 (October 2017): 154–69. 10.1016/j.rse.2017.07.029.

Claar, Danielle C., Lisa Szostek, Jamie M. McDevitt-Irwin, Julian J. Schanze, and Julia K. Baum. 2018. ‘Global Patterns and Impacts of El Niño Events on Coral Reefs: A Meta-Analysis’. PLOS ONE 13 (2): e0190957. 10.1371/journal.pone.0190957.

Colgan, Mitchell W. 1987. ‘Coral Reef Recovery on Guam (Micronesia) After Catastrophic Predation by Acanthaster Planci’. Ecology 68 (6): 1592–605. 10.2307/1939851.

Connell, Joseph H. 1973. ‘Population Ecology of Reef-Building Corals’. In Biology and Geology of Coral Reefs, 205–45. Elsevier. 10.1016/B978-0-12-395526-5.50015-8.

Connell, Joseph H., Terence P. Hughes, and Carden C. Wallace. 1997. ‘A 30-Year Study of Coral Abundance, Recruitment, and Disturbance at Several Scales in Space and Time’. Ecological Monographs 67 (4): 461–88. 10.1890/0012-9615(1997)067%5B0461:AYSOCA%5D2.0.CO;2.

DeMerlis, Allyson, Amanda Kirkland, Madeline L. Kaufman, Anderson B. Mayfield, Nathan Formel, Graham Kolodziej, Derek P. Manzello, Diego Lirman, Nikki Traylor-Knowles, and Ian C. Enochs. 2022. ‘Pre-Exposure to a Variable Temperature Treatment Improves the Response of Acropora Cervicornis to Acute Thermal Stress’. Coral Reefs 41 (2): 435–45. 10.1007/s00338-022-02232-z.

Dixon, Adele M., Piers M. Forster, Scott F. Heron, Anne M. K. Stoner, and Maria Beger. 2022. ‘Future Loss of Local-Scale Thermal Refugia in Coral Reef Ecosystems’. PLOS Climate 1 (2): e0000004. 10.1371/journal.pclm.0000004.

Donovan, Mary K., Thomas C. Adam, Andrew A. Shantz, Kelly E. Speare, Katrina S. Munsterman, Mallory M. Rice, Russell J. Schmitt, Sally J. Holbrook, and Deron E. Burkepile. 2020. ‘Nitrogen Pollution Interacts with Heat Stress to Increase Coral Bleaching across the Seascape’. Proceedings of the National Academy of Sciences 117 (10): 5351–57. 10.1073/pnas.1915395117.

Edmunds, Peter, Adrian Cheh, and Scott Burgess. 2025. ‘A Physiological Crisis Drives the Coral Recruitment Bottleneck’. Biology Letters 21 (6): 20250103. 10.1098/rsbl.2025.0103.

Ferrara, Erik F., Anna Roik, Franziska Wöhrmann-Zipf, and Maren Ziegler. 2024. ‘Thermal Preconditioning Modulates Coral Physiology and Heat Tolerance: A Multi-Species Perspective’. bioRxiv, ahead of print, 2024. 10.1101/2024.07.18.604102.

García, Francisca C., Eslam O. Osman, Neus Garcias-Bonet, Nathalia Delgadillo-Ordoñez, Erika P. Santoro, Inês Raimundo, Helena D. M. Villela, Christian R. Voolstra, and Raquel S. Peixoto. 2024. ‘Seasonal Changes in Coral Thermal Threshold Suggest Species-Specific Strategies for Coping with Temperature Variations’. Communications Biology 7 (1): 1680. 10.1038/s42003-024-07340-w.

Glynn, Victoria M., Laura Fernandes De Barros Marangoni, Maxime Guglielmetti, Eunice R. Tapia, Viviane Ali, Helio Quintero, E. Catalina Rodriguez Guerra, et al. 2025. ‘The Role of Holobiont Composition and Environmental History in Thermotolerance of Tropical Eastern Pacific Corals’. Current Biology 35 (13): 3048–3063.e7. 10.1016/j.cub.2025.05.035.

Graham, N. A. J., K. L. Nash, and J. T. Kool. 2011. ‘Coral Reef Recovery Dynamics in a Changing World’. Coral Reefs 30 (2): 283–94. 10.1007/s00338-010-0717-z.

Graham, Nicholas A. J., Shaun K. Wilson, Cassandra E. Benkwitt, Rodney Bonne, Rodney Govinden, and James P. W. Robinson. 2024. ‘Increased Resilience and a Regime Shift Reversal through Repeat Mass Coral Bleaching’. Ecology Letters 27 (12): e14454. 10.1111/ele.14454.

Grigg, Richard W. 1983. ‘Community Structure, Succession and Development of Coral Reefs in Hawaii’. Marine Ecology Progress Series 11 (1983): 1–14.

Grottoli, Andréa G., Lisa J. Rodrigues, and James E. Palardy. 2006. ‘Heterotrophic Plasticity and Resilience in Bleached Corals’. Nature 440 (7088): 1186–89. 10.1038/nature04565.

Grottoli, Andréa G., Mark E. Warner, Stephen J. Levas, Matthew D. Aschaffenburg, Verena Schoepf, Michael McGinley, Justin Baumann, and Yohei Matsui. 2014. ‘The Cumulative Impact of Annual Coral Bleaching Can Turn Some Coral Species Winners into Losers’. Global Change Biology 20 (12): 3823–33. 10.1111/gcb.12658.

Hackerott, Serena, Harmony A. Martell, and Jose M. Eirin-Lopez. 2021. ‘Coral Environmental Memory: Causes, Mechanisms, and Consequences for Future Reefs’. Trends in Ecology & Evolution 36 (11): 1011–23. 10.1016/j.tree.2021.06.014.

Hall, V. R., and T. P. Hughes. 1996. ‘Reproductive Strategies of Modular Organisms: Comparative Studies of Reef-Building Corals’. Ecology 77 (3): 950–63. 10.2307/2265514.

Hill, Jos, and CLIVE Wilkinson. 2004. ‘Methods for Ecological Monitoring of Coral Reefs’. Australian Institute of Marine Science, Townsville 117 (2004).

Hoegh-Guldberg, Ove. 1999. ‘Climate Change, Coral Bleaching and the Future of the World’s Coral Reefs’. Marine and Freshwater Research, ahead of print, 1999. 10.1071/MF99078.

Holbrook, Neil J., Hillary A. Scannell, Alexander Sen Gupta, Jessica A. Benthuysen, Ming Feng, Eric C. J. Oliver, Lisa V. Alexander, et al. 2019. ‘A Global Assessment of Marine Heatwaves and Their Drivers’. Nature Communications 10 (1): 2624. 10.1038/s41467-019-10206-z.

Holbrook, Sally J., Thomas C. Adam, Peter J. Edmunds, Russell J. Schmitt, Robert C. Carpenter, Andrew J. Brooks, Hunter S. Lenihan, and Cheryl J. Briggs. 2018. ‘Recruitment Drives Spatial Variation in Recovery Rates of Resilient Coral Reefs’. Scientific Reports 8 (1): 7338. 10.1038/s41598-018-25414-8.

Howells, Emily J., Ray Berkelmans, Madeleine J. H. Van Oppen, Bette L. Willis, and Line K. Bay. 2013. ‘Historical Thermal Regimes Define Limits to Coral Acclimatization’. Ecology 94 (5): 1078–88. 10.1890/12-1257.1.

Huffmyer, Ariana S., Emma L. Strand, Serena Hackerott, Kevin H. Wong, Danielle M. Becker, Dennis Conetta, Kristina X. Terpis, et al. 2025. ‘Seasonal Plasticity of Symbiotic Strategies Clarifies Coral Holobiont Resistance and Resilience’. bioRxiv: The Preprint Server for Biology, ahead of print, 2025. 10.1101/2025.11.20.689604.

Hughes, Terry P., Kristen D. Anderson, Sean R. Connolly, Scott F. Heron, James T. Kerry, Janice M. Lough, Andrew H. Baird, et al. 2018. ‘Spatial and Temporal Patterns of Mass Bleaching of Corals in the Anthropocene’. Science 359 (6371): 80–83. 10.1126/science.aan8048.

Hughes, Terry P., James T. Kerry, Mariana Álvarez-Noriega, Jorge G. Álvarez-Romero, Kristen D. Anderson, Andrew H. Baird, Russell C. Babcock, et al. 2017. ‘Global Warming and Recurrent Mass Bleaching of Corals’. Nature 543 (7645): 373–77. 10.1038/nature21707.

Hughes, Terry P., James T. Kerry, Andrew H. Baird, Sean R. Connolly, Tory J. Chase, Andreas Dietzel, Tessa Hill, et al. 2019. ‘Global Warming Impairs Stock–Recruitment Dynamics of Corals’. Nature 568 (7752): 387–90. 10.1038/s41586-019-1081-y.

Humanes, Adriana, Liam Lachs, Elizabeth A. Beauchamp, John C. Bythell, Alasdair J. Edwards, Yimnang Golbuu, Helios M. Martinez, et al. 2022. ‘Within-Population Variability in Coral Heat Tolerance Indicates Climate Adaptation Potential’. Proceedings of the Royal Society B: Biological Sciences 289 (1981): 20220872. 10.1098/rspb.2022.0872.

James, Anna K., Libe Washburn, Chris Gotschalk, Stéphane Maritorena, Alice Alldredge, Craig E. Nelson, James L. Hench, James J. Leichter, Alex S. J. Wyatt, and Craig A. Carlson. 2020. ‘An Island Mass Effect Resolved Near Mo’orea, French Polynesia’. Frontiers in Marine Science 7 (January 2020): 16. 10.3389/fmars.2020.00016.

Johns, Kerryn A., Michael J. Emslie, Andrew S. Hoey, Kate Osborne, Michelle J. Jonker, and Alistair J. Cheal. 2018. ‘Macroalgal Feedbacks and Substrate Properties Maintain a Coral Reef Regime Shift’. Ecosphere 9 (7): e02349. 10.1002/ecs2.2349.

Johnston, Erika C., Chelsie W. W. Counsell, Tayler L. Sale, Scott C. Burgess, and Robert J. Toonen. 2020. ‘The Legacy of Stress: Coral Bleaching Impacts Reproduction Years Later’. Functional Ecology 34 (11): 2315–25. 10.1111/1365-2435.13653.

Kay, Matthew. 2023. Tidybayes: Tidy Data and Geoms for Bayesian Models. 10.5281/zenodo.1308151.

Kinzie, Robert A., Michelle Takayama, Scott R. Santos, and Mary Alice Coffroth. 2001. ‘The Adaptive Bleaching Hypothesis: Experimental Tests of Critical Assumptions’. The Biological Bulletin 200 (1): 51–58. 10.2307/1543084.

Kopecky, Kai L., Sally J. Holbrook, Emalia Partlow, Madeline Cunningham, and Russell J. Schmitt. 2024. ‘Changing Disturbance Regimes, Material Legacies, and Stabilizing Feedbacks: Dead Coral Skeletons Impair Key Recovery Processes Following Coral Bleaching’. Global Change Biology 30 (9): e17504. 10.1111/gcb.17504.

Leinbach, Sarah E., Kelly E. Speare, Ashley M. Rossin, Daniel M. Holstein, and Marie E. Strader. 2021. ‘Energetic and Reproductive Costs of Coral Recovery in Divergent Bleaching Responses’. Scientific Reports 11 (1): 23546. 10.1038/s41598-021-02807-w.

Loya, Y., K. Sakai, K. Yamazato, Y. Nakano, H. Sambali, and R. Van Woesik. 2001. ‘Coral Bleaching: The Winners and the Losers’. Ecology Letters 4 (2): 122–31. 10.1046/j.1461-0248.2001.00203.x.

Lüdecke, Daniel, Mattan S. Ben-Shachar, Indrajeet Patil, Philip Waggoner, and Dominique Makowski. 2021. ‘performance: An R Package for Assessment, Comparison and Testing of Statistical Models’. Journal of Open Source Software 6 (60): 3139. 10.21105/joss.03139.

Madin, Joshua S., Andrew H. Baird, Maria Dornelas, and Sean R. Connolly. 2014. ‘Mechanical Vulnerability Explains Size-dependent Mortality of Reef Corals’. Ecology Letters 17 (8): 1008–15. 10.1111/ele.12306.

Maher, Rebecca L., Emily R. Schmeltzer, Sonora Meiling, Ryan McMinds, Leïla Ezzat, Andrew A. Shantz, Thomas C. Adam, et al. 2020. ‘Coral Microbiomes Demonstrate Flexibility and Resilience Through a Reduction in Community Diversity Following a Thermal Stress Event’. Frontiers in Ecology and Evolution 8 (October 2020): 555698. 10.3389/fevo.2020.555698.

McClanahan, T. R., and Nyawira A. Muthiga. 2021. ‘Oceanic Patterns of Thermal Stress and Coral Community Degradation on the Island of Mauritius’. Coral Reefs 40 (1): 53–74. 10.1007/s00338-020-02015-4.

Morais, Juliano, Sterling B. Tebbett, Renato A. Morais, and David R. Bellwood. 2024. ‘Natural Recovery of Corals after Severe Disturbance’. Ecology Letters 27 (1): e14332. 10.1111/ele.14332.

Moritz, C., Sj Brandl, H. Rouzé, J. Vii, G. Pérez-Rosales, P. Bosserelle, Y. Chancerelle, et al. 2021. ‘Long-Term Monitoring of Benthic Communities Reveals Spatial Determinants of Disturbance and Recovery Dynamics on Coral Reefs’. Marine Ecology Progress Series 672 (August 2021): 141–52. 10.3354/meps13807.

Muscatine, L., L. R. McCloskey, and R. E. Marian. 1981. ‘Estimating the Daily Contribution of Carbon from Zooxanthellae to Coral Animal Respiration1’. Limnology and Oceanography 26 (4): 601–11. 10.4319/lo.1981.26.4.0601.

Nakamura, T., and R. Van Woesik. 2001. ‘Water-Flow Rates and Passive Diffusion Partially Explain Differential Survival of Corals during the 1998 Bleaching Event’. Marine Ecology Progress Series 212 (2001): 301–4. 10.3354/meps212301.

Naugle, Melissa S., Hugo Denis, Véronique J. L. Mocellin, Patrick W. Laffy, Iva Popovic, Line K. Bay, and Emily J. Howells. 2024. ‘Heat Tolerance Varies Considerably within a Reef-Building Coral Species on the Great Barrier Reef’. Communications Earth & Environment 5 (1): 525. 10.1038/s43247-024-01649-4.

Oliver, Eric C. J., Michael T. Burrows, Markus G. Donat, Alex Sen Gupta, Lisa V. Alexander, Sarah E. Perkins-Kirkpatrick, Jessica A. Benthuysen, et al. 2019. ‘Projected Marine Heatwaves in the 21st Century and the Potential for Ecological Impact’. Frontiers in Marine Science 6 (December 2019): 734. 10.3389/fmars.2019.00734.

Oxenford, HA, R. Roach, and Angelique Brathwaite. 2008. ‘Large Scale Coral Mortality in Barbados: A Delayed Response to the 2005 Bleaching Episode’. Proceedings of the 11th International Coral Reef Symposium Ft. Lauderdale: Florida, 2008, 7–11.

Pavoni, Gaia, Massimiliano Corsini, Federico Ponchio, Alessandro Muntoni, Clinton Edwards, Nicole Pedersen, Stuart Sandin, and Paolo Cignoni. 2022. ‘TagLab: AI-Assisted Annotation for the Fast and Accurate Semantic Segmentation of Coral Reef Orthoimages’. Journal of Field Robotics 39 (3): 246–62. 10.1002/rob.22049.

Pratchett, Morgan S., Dominique McCowan, Jeffrey A. Maynard, and Scott F. Heron. 2013. ‘Changes in Bleaching Susceptibility among Corals Subject to Ocean Warming and Recurrent Bleaching in Moorea, French Polynesia’. PLoS ONE 8 (7): e70443. 10.1371/journal.pone.0070443.

Putnam, Hollie M., Michael Stat, Xavier Pochon, and Ruth D. Gates. 2012. ‘Endosymbiotic Flexibility Associates with Environmental Sensitivity in Scleractinian Corals’. Proceedings of the Royal Society B: Biological Sciences 279 (1746): 4352–61. 10.1098/rspb.2012.1454.

R Core Team. 2024. R: A Language and Environment for Statistical Computing. Manual. Vienna, Austria: R Foundation for Statistical Computing. https://www.R-project.org/.

Rasher, Douglas B., and Mark E. Hay. 2010. ‘Seaweed Allelopathy Degrades the Resilience and Function of Coral Reefs’. Communicative & Integrative Biology 3 (6): 564–66. 10.4161/cib.3.6.12978.

Reimer, James Davis, Raquel S. Peixoto, Sarah W. Davies, Nikki Traylor-Knowles, Morgan L. Short, Rafael A. Cabral-Tena, John A. Burt, et al. 2024. ‘The Fourth Global Coral Bleaching Event: Where Do We Go from Here?’ Coral Reefs 43 (4): 1121–25. 10.1007/s00338-024-02504-w.

Robinson, James P. W., Shaun K. Wilson, and Nicholas A. J. Graham. 2019. ‘Abiotic and Biotic Controls on Coral Recovery 16 Years after Mass Bleaching’. Coral Reefs 38 (6): 1255–65. 10.1007/s00338-019-01831-7.

Rowan, Rob. 2004. ‘Thermal Adaptation in Reef Coral Symbionts’. Nature 430 (7001): 742–742. 10.1038/430742a.

Sing Wong, Amy, Spyridon Vrontos, and Michelle L. Taylor. 2022. ‘An Assessment of People Living by Coral Reefs over Space and Time’. Global Change Biology 28 (23): 7139–53. 10.1111/gcb.16391.

Skirving, William, Benjamin Marsh, Jacqueline De La Cour, Gang Liu, Andy Harris, Eileen Maturi, Erick Geiger, and C. Mark Eakin. 2020. ‘CoralTemp and the Coral Reef Watch Coral Bleaching Heat Stress Product Suite Version 3.1’. Remote Sensing 12 (23): 3856. 10.3390/rs12233856.

SNO CORAIL. 2025. ‘Temporal Physical, Physicochemical, and Biological Variability in Pacific Coral Reef Ecosystems Monitored by the SNO Corail’. Version 2025. SEANOE. 10.17882/108741.

Sommer, Brigitte, Jessica M. Hodge, Liam Lachs, James Cant, John M. Pandolfi, and Maria Beger. 2024. ‘Decadal Demographic Shifts and Size-Dependent Disturbance Responses of Corals in a Subtropical Warming Hotspot’. Scientific Reports 14 (1): 6327. 10.1038/s41598-024-56890-w.

Spady, Blake L., William J. Skirving, Erick F. Geiger, Neal E. Cantin, Gang Liu, Jacqueline L. De La Cour, Peter J. Mumby, Andrew Norrie, and Derek P. Manzello. 2025. ‘Satellite-Based Analysis of an Unverified Mass Coral Bleaching Event on the Great Barrier Reef in 2021’. Coral Reefs 44 (4): 1275–85. 10.1007/s00338-025-02690-1.

Speare, Kelly E., Thomas C. Adam, Erin M. Winslow, Hunter S. Lenihan, and Deron E. Burkepile. 2022. ‘Size-dependent Mortality of Corals during Marine Heatwave Erodes Recovery Capacity of a Coral Reef’. Global Change Biology 28 (4): 1342–58. 10.1111/gcb.16000.

Sully, S., D. E. Burkepile, M. K. Donovan, G. Hodgson, and R. Van Woesik. 2019. ‘A Global Analysis of Coral Bleaching over the Past Two Decades’. Nature Communications 10 (1): 1264. 10.1038/s41467-019-09238-2.

Thomas, Luke, Noah H. Rose, Rachael A. Bay, Elora H. López, Megan K. Morikawa, Lupita Ruiz-Jones, and Stephen R. Palumbi. 2018. ‘Mechanisms of Thermal Tolerance in Reef-Building Corals across a Fine-Grained Environmental Mosaic: Lessons from Ofu, American Samoa’. Frontiers in Marine Science 4 (February 2018): 434. 10.3389/fmars.2017.00434.

Van Oppen, Madeleine J. H., Pim Bongaerts, Pedro Frade, Lesa M. Peplow, Sarah E. Boyd, Hieu T. Nim, and Line K. Bay. 2018. ‘Adaptation to Reef Habitats through Selection on the Coral Animal and Its Associated Microbiome’. Molecular Ecology 27 (14): 2956–71. 10.1111/mec.14763.

Van Oppen, Madeleine J. H., James K. Oliver, Hollie M. Putnam, and Ruth D. Gates. 2015. ‘Building Coral Reef Resilience through Assisted Evolution’. Proceedings of the National Academy of Sciences 112 (8): 2307–13. 10.1073/pnas.1422301112.

Van Woesik, R., A. Irikawa, R. Anzai, and T. Nakamura. 2012. ‘Effects of Coral Colony Morphologies on Mass Transfer and Susceptibility to Thermal Stress’. Coral Reefs 31 (3): 633–39. 10.1007/s00338-012-0911-2.

Van Woesik, R., K. Sakai, A. Ganase, and Y. Loya. 2011. ‘Revisiting the Winners and the Losers a Decade after Coral Bleaching’. Marine Ecology Progress Series 434 (July 2011): 67–76. 10.3354/meps09203.

Vermeij, Mark J. A., and Stuart A. Sandin. 2008. ‘Density-Dependent Settlement and Mortality Structure the Earliest Life Phases of a Coral Population’. Ecology 89 (7): 1994–2004. 10.1890/07-1296.1.

Vompe, Alex D., Hannah E. Epstein, Kelly E. Speare, Emily R. Schmeltzer, Thomas C. Adam, Deron E. Burkepile, Thomas J. Sharpton, and Rebecca Vega Thurber. 2024. ‘Microbiome Ecological Memory and Responses to Repeated Marine Heatwaves Clarify Variation in Coral Bleaching and Mortality’. Global Change Biology 30 (1): e17088. 10.1111/gcb.17088.

Voolstra, Christian R., and Maren Ziegler. 2020. ‘Adapting with Microbial Help: Microbiome Flexibility Facilitates Rapid Responses to Environmental Change’. BioEssays 42 (7): 2000004. 10.1002/bies.202000004.

Winslow, Erin M., Kelly E. Speare, Thomas C. Adam, Deron E. Burkepile, James L. Hench, and Hunter S. Lenihan. 2024. ‘Corals Survive Severe Bleaching Event in Refuges Related to Taxa, Colony Size, and Water Depth’. Scientific Reports 14 (1): 9006. 10.1038/s41598-024-58980-1.

Wyatt, Alex S. J., James J. Leichter, Libe Washburn, Li Kui, Peter J. Edmunds, and Scott C. Burgess. 2023. ‘Hidden Heatwaves and Severe Coral Bleaching Linked to Mesoscale Eddies and Thermocline Dynamics’. Nature Communications 14 (1): 25. 10.1038/s41467-022-35550-5.

Ziegler, Maren, Carsten G. B. Grupstra, Marcelle M. Barreto, Martin Eaton, Jaafar BaOmar, Khalid Zubier, Abdulmohsin Al-Sofyani, Adnan J. Turki, Rupert Ormond, and Christian R. Voolstra. 2019. ‘Coral Bacterial Community Structure Responds to Environmental Change in a Host-Specific Manner’. Nature Communications 10 (1): 3092. 10.1038/s41467-019-10969-5.

## References

Spady, B. L., Skirving, W. J., Geiger, E. F., Cantin, N. E., Liu, G., De La Cour, J. L., Mumby, P. J., Norrie, A., & Manzello, D. P. (2025). Satellite-based analysis of an unverified mass coral bleaching event on the Great Barrier Reef in 2021. Coral Reefs, 44(4), 1275–1285. 10.1007/s00338-025-02690-1

